# Eye in a Disk: eyeIntegration human pan-eye and body transcriptome database version 1.0

**DOI:** 10.1101/579482

**Authors:** Vinay Swamy, David McGaughey

**Affiliations:** Ophthalmic Genetics and Visual Function Branch, National Eye Institute, National Institutes of Health

## Abstract

**PURPOSE:** To develop an accessible and reliable RNA-seq transcriptome database of healthy human eye tissues and a matching reactive web application to query gene expression in eye and body tissues. METHODS: We downloaded the raw sequnce data for 1375 RNA-seq samples across 54 tissues in the GTEx project as a non-eye reference set.

We then queried several public repositories to find all healthy, non-perturbed, human eye-related tissue RNA-seq samples. The 916 eye and 1375 GTEx samples were sent into a Snakemake-based reproducible pipeline we wrote to quantify all known transcripts and genes, removes samples with poor sequence quality and mislabels, normalizes expression values across each tissue, performs 882 differential expression tests, calculates GO term enrichment, and outputs all as a single SQLite database file: the Eye in a Disk (EiaD) dataset. Furthermore, we rewrote the web application eyeIntegration (https://eyeIntegration.nei.nih.gov) to display EiaD.

**RESULTS:** The new eyeIntegration portal provides quick visualization of human eye-related transcriptomes published to date by database version, gene/transcript, 19 eye tissues, and 54 body tissues. As a test of the value of this unified pan-eye dataset, we showed that fetal and organoid retina are highly similar at a pan-transcriptome level but display distinct differences in certain pathways and gene families like protocadherin and HOXB family members.

**CONCLUSION:** The eyeIntegration v1.0 web app serves the pan-human eye and body transcriptome dataset, EiaD. This offers the eye community a powerful and quick means to test hypotheses on human gene and transcript expression across 54 body and 19 eye tissues.

## Introduction

### RNA-seq is the predominant technology for deciphering transcriptomes

From anterior to posterior along the light trajectory, the human eye is composed of the cornea, lens, retina, retinal pigment epithelium (RPE), and choroid. The differentiation, maturation, and function of these tissues is mediated through spatial and temporal specific transcript and gene expression patterns, also known as the transcriptome. Today, RNA-sequencing (RNA-seq) is the predominant technology for quantifying the transcriptome. Analysis of the transcripts’ expression across tissue, time, and perturbation allows researchers to decipher the genetic controls of eye development and function. To this end, a wide variety of human tissue sources have been used to assess gene function, including primary tissue (fetal and post-mortem), differentiated stem cells, immortalized cell lines, and most recently, organoids. These tissue types have been deeply sequenced across the cornea^1–7^, lens^8^, retina^9–17^, and RPE (choroid)^14, 17–34^.

### The GTEx gene expression web app lacks eye-specific tissues

The Genotype-Tissue Expression (GTEx) Project has generated RNA-seq data across dozens of post-mortem human tissues from hundreds of unique donors and presents the gene and transcript level data in a comprehensive and user-friendly web app (https://gtexportal.org/); however eye tissues have not been included^35, 36^. Recently Ratnapriya et al. have published a huge set of post-mortem retina, both normal and with varying degrees of age-related macular degeneration (AMD) and the GTEx project is providing the data as a download link. This data, as of June 2019, is not available in the interactive GTEx visualizations^37^. The Sequence Read Archive (SRA) and European Nucleotide Archive (ENA) are the primary repositories for all raw sequence data and two groups have quantified large portions of the RNA-seq data, including some human eye tissues, from the SRA: recount2 and ARCHS4^38, 39^. To date, no curation of the sample level metadata has been done, therefore it is challenging to parse out which eye tissues are present and even more difficult to determine whether any samples were chemically or genetically perturbed. More targeted web resources that allow researchers to quickly assess gene expression in eye tissues include iSYTE, EXPRESS, and retina.Tigem.it^16, 40, 41^. However iSYTE only includes lens samples, EXPRESS is limited to a subset of mouse lens and retina samples, and retina.Tigem.it is retina only. We thus aimed our efforts at developing an easily accessible and reliable RNA-seq based transcriptome database of healthy human eye tissues and a matching reactive web application to query gene expression in eye and body tissues.

### The eyeIntegration app interactively serves huge GTEx and human eye tissue datasets (EiaD)

The eyeIntegration web resource (https://eyeIntegration.nei.nih), originally released in 2017 at version 0.6, provides the largest set of transcriptomes from hand-curated human eye tissues along with hundreds of GTEx tissue samples^42^. This interactive web app allows for quick transcript and gene comparisons across many eye tissues and dozens of other body tissues. The dataset that the original eyeIntegration web app served was created with a series of scripts, several of which were run interactively to manually assess quality control for the samples. The interactive nature of some of the steps precluded efficient and regular data updates for the data.

To better meet the needs of the eye research community we have re-written the bioinformatic pipeline that creates the eye and body RNA-seq dataset to allow for regular, versioned updates for eyeIntegration. We call this reproducible and versioned transcriptome dataset “Eye in a Disk” (EiaD). The pipeline automates the EiaD creation, ensures full reproducibility of the results, allow for external data comparison, provides consistent sample quality control, and improves efficiency for future sample updates. The 2019 EiaD dataset contains several new tissue types, full gene product quantification, along with hundreds of new samples and improved sample labeling. The eyeIntegration web app has also been re-written to provide many new features, including versioned EiaD datasets, custom URL shortcut creation, new visualizations, improved data table searching, easy download of core datasets, and local install of the entire interactive resource with three commands. Additionally, we are prototyping new tools to display single cell RNA-seq (scRNA-seq) data to provide researchers access to cell type specific information about gene expression across murine retinal development.

### The EiaD dataset can be used to identify potential avenues to improve retina organoid maturation

Retina organoids are an increasingly popular means to model human retina development. We used our pan-study EiaD dataset to show that, at a pan-transcriptome level, organoids are highly similar to early fetal retina tissue. We also show that important temporal gene expression patterns in the fetal retina tissue are recapitulated in the organoids. As the organoid differentiation methods do not yet produce fully mature retina, we focused on identifying differentially expressed processes between organoid retina and embryonic retina and detected, for example, identifying protocadherin and HOXB family gene expression differences which suggest targetable pathways to improve and benchmark organoid differentiation methods.

## Methods

### Identification of potential eye samples

We exhaustively searched the SRA with the SRAdb R package for eye related tissues using the query ‘cornea|retina|RPE|macula|fovea|choroid|sclera|iris|lens|eye’ across all columns and rows in the ‘SRA’ table^43, 44^. As the SRAdb is being deprecated we also ran searches on the SRA and Gene Expression Omnibus (GEO) web pages with as follows: ((“Homo sapiens”[orgn: txid9606]) AND (transcriptomic[Source]) AND (“2019/01/01”[Publication Date] : “3000”[Publication Date]) AND (retina[Text Word] OR RPE[Text Word] OR macula[Text Word] OR fovea[Text Word] OR choroid[Text Word] OR sclera[Text Word] OR iris[Text Word] OR lens[Text Word] OR cornea[Text Word] OR ‘trabecular meshwork’[Text Word] OR ‘canals of schlemm’[Text Word] OR ‘cillary body’[Text Word] OR ‘optic nerve’[Text Word] OR ‘laminar cribosa’[Text Word] OR retina[Title] OR RPE[Title] OR macula[Title] OR fovea[Title] OR choroid[Title] OR sclera[Title] OR iris[Title] OR lens[Title] OR cornea[Title] OR ‘trabecular meshwork’[Title] OR ‘canals of schlemm’[Title] OR ‘cillary body’[Title] OR ‘optic nerve’[Title] OR ‘laminar cribosa’[Title]))”. We hand selected relevant studies and selected healthy, control or unmodified samples spanning primary adult tissue, primary fetal tissue, induced pluripotent stem cell (iPSC)-derived tissue, stem cell derived organoids, and immortalized cell lines. In order to compare gene expression in the eye against expression in other body tissues, we obtained samples from 54 different body tissues from the GTEx project. Using SRA metadata from each study we extracted sample and run accessions, library type, tissue of origin, and sub-tissue of origin. Any of the preceding information missing from the SRA metadata was added by hand, when available. Stem cell-derived tissues and cell lines are marked as sub-tissues of the tissue they model.

### Raw data download and quantification

We downloaded the relevant SRA files for each sample directly from the NCBI ftp server using the file transfer software Aspera. SRA files were converted to FASTQ format using the tool fastq-dump from the SRAtoolkit software package^43^. Samples only available in the BAM format were converted to FASTQ format using SAMTools^45^. Sample transcriptomes were quantified using the alignment free quantification software Salmon, using transcriptomic index built from gencode v28 protein coding transcript sequences using the transcriptomic aligner Salmon^46, 47^. Using the resulting expression quantification, we identified lowly or unused transcripts within the gencode annotation, and removed transcripts that accounted for 5% or less of the total expression for its parent gene as per Sonneson et al^48^. Samples were re-quantified against a transcriptomic index built on the filtered transcript sequences. The Salmon count values were quantified as (transcript) length scaled Transcripts Per Million (TPM) to the transcript and gene level using tximport^49^.

### Quality control

We first removed samples with a Salmon calculated mapping rate less than 40%. This value was selected as being the far left tail of the distribution of mapping rates across samples (Supplemental Figure 2). We removed lowly expressed genes by calculating the median expression across all samples for each gene and kept genes that had a median count >200 across all samples. To reduce the noise from experimental variability between each study, we normalized samples by sequence library size using the calcNormFactors function from the edgeR R package, and then quantile smoothed expression data using the R package qsmooth at the tissue level^50, 51^. In a change from our previous eyeIntegration work^42^, we now correct our counts for mapping rate and tissue type with the limma batchEffects function^52^. The transformed values are used for the box plot and t-SNE visualizations.

To identify outliers we followed an approach similar to a method in Wright et al^53^. Briefly, we first selected the 3000 genes with the highest variance across all samples and then for each sub-tissue type *T*, and each sample *i* in *T*, we first calculated *r_i_*, the average correlation between *i* and all other samples in *T*. Next, we calculated *D_i_*, where 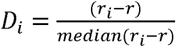 and *r*. is the grand mean of all *r_i_* for *i* in *T*,. We removed samples with *D_i_* < −17.5; we determined this threshold by generating a tSNE plot of our samples, and visually identifying outliers in adult retina tissue. The(*max*(*D_i_*)) amongst these outliers was −17.58 and from this we chose −17.5 as our outlier threshold.

To calculate pearson correlation (*R*^2^) between GTEx-calculated TPM gene values and our GTEx TPM gene values, we downloaded “GTEx_Analysis_2016-01-15_v7_RNASeQCv1.1.8_gene_tpm.gct.gz” and matched against our GTEx TPM values, running the pearson correlation with fo’a2(l,bW + 0.01) values as per Zhang et al. with the cor function in R^54^.

### Differential Gene Expression Analysis and GO term enrichment

We used the non-transformed length scaled TPM values to determine differential gene expression between different sub-tissue types. First, we generated a synthetic body set to serve as single representative sub-tissue type for pan-body gene expression by randomly sampling GTEx tissues. We used the voom function from the limma R package to convert gene expression to precision weights, and then performed pairwise differential expression tests for all combinations of eye sub-tissues (using mapping rate as a covariate), the synthetic body tissue, and human body tissues using an empirical Bayes test^52, 55^. We extracted significant genes (FDR p < 0.01) for all 882 comparisons and used these to calculate GO enrichment. The significant gene list for each eye sub-tissue was split into upregulated and down regulated sets and each set was tested for enrichment using the enrichGO function from the clusterProfiler R package (q-value < 0.01)^56^.

### eyeIntegration web app and R package

The data generated in the above steps is consolidated into a SQLite database, with the original dataset for eyeIntegration and the new 2019 EiaD dataset each getting a separate database file. The code that creates the eyeIntegration web app is written in Shiny and R and has been wrapped into an R package (https://github.com/davemcg/eyeIntegration_app/) that can be deployed on a local computer or a web server (https://eyeIntegration.nei.nih.gov). The app can be deployed on a local computer with 50GB of free disk space by running three commands in R: “devtools::install_github(‘davemcg/eyeIntegration_app’)”, “eyeIntegrationApp::get_eyeIntegration_datasets()”, and “eyeIntegrationApp::run_eyeIntegration()”.

### Snakemake reproducible pipeline

While the sample search and metadata parsing in a semi-curated process, the processing from the raw data to the creation of the SQLite EiaD database underlying eyeIntegration is wrapped in a Snakemake pipeline, which ensures full reproducibility of the results^57^. We make the code for the pipeline available at https://github.com/davemcg/EiaD_build.

### scRNA-seq processing

The eyeIntegration site, as of June 2019, hosts two large scRNA-seq datasets from Macosko et al. and Clark et al^58, 59^. We use the processed gene count data directly from each group, as well as their cluster assignments which specify what cell type each individual cell is. The count data is mean averaged to the cell type, age, and gene level for the single cell expression section of eyeIntegration. We also display t-SNE and UMAP-based two-dimensional visualizations of the Macosko and Clark data, respectively, in the web app. For detail so the t-SNE processing we did on the Macosko dataset, see the methods of Bryan et al^42^.

### Power Calculation

We use the ssizeRNA R package to calculate power (p) across samples (n) at an FDR of 0.05^60^. Important parameters for ssize RNA include the variability (dispersions for the samples and genes), which were calculated directly from our EiaD length scaled TPM values by the edgeR packages estimateCommonDisp and estimateTagwiseDisp. The code to calculate the power is given as ‘power_calc.R’.

### Manuscript as code and reproducibility

This manuscript’s figures, tables, and most numbers, are all created and laid out in a R markdown document that interweaves code and text. The knitr and pandoc program is used to lay out the figures and tables an output a docx file. The code that generates this manuscript can be found at https://github.com/davemcg/eyeIntegration_v1_app_manuscript.

The relevant code-bases (https://github.com/davemcg/eyeIntegration_v1_app_manuscript, https://github.com/davemcg/EiaD_build) and the EiaD dataset itself has been deposited into Zenodo with accessio 10.5281/zenodo.3238677 to ensure the data can be accessed in the future, even should eyeIntegration and GitHu become inaccessible in the future.

## Results

### EiaD 2019 contains 24 new human eye RNA-seq studies, 448 new Retina AMD samples, 207 new eye samples, and 16 total eye sub-tissue types

Our query on May 8th 2019 to the SRA found 107 potentially relevant studies. We removed non-pertinent studies and selected healthy or unmodified tissue from each relevant study for a total, including of 46 studies, 30 of which are new to the 2019 EiaD dataset. The 2019 EiaD dataset contains 835 human eye tissue samples and also includes 1314 GTEx samples across 54 tissues for easy comparison (Table 1, Supplemental Table 1). The 2019 EiaD contains 6 undifferentiated iPSC, 56 cornea, 4 lens, 648 retina, and 121 RPE (choroid) samples; in total we have added 655 new samples to the 2019 EiaD (Figure 1). We refer to native-tissue extracted RPE as RPE (choroid) because it is not possible to remove the choroid from the RPE without culturing.

**Figure 1:**
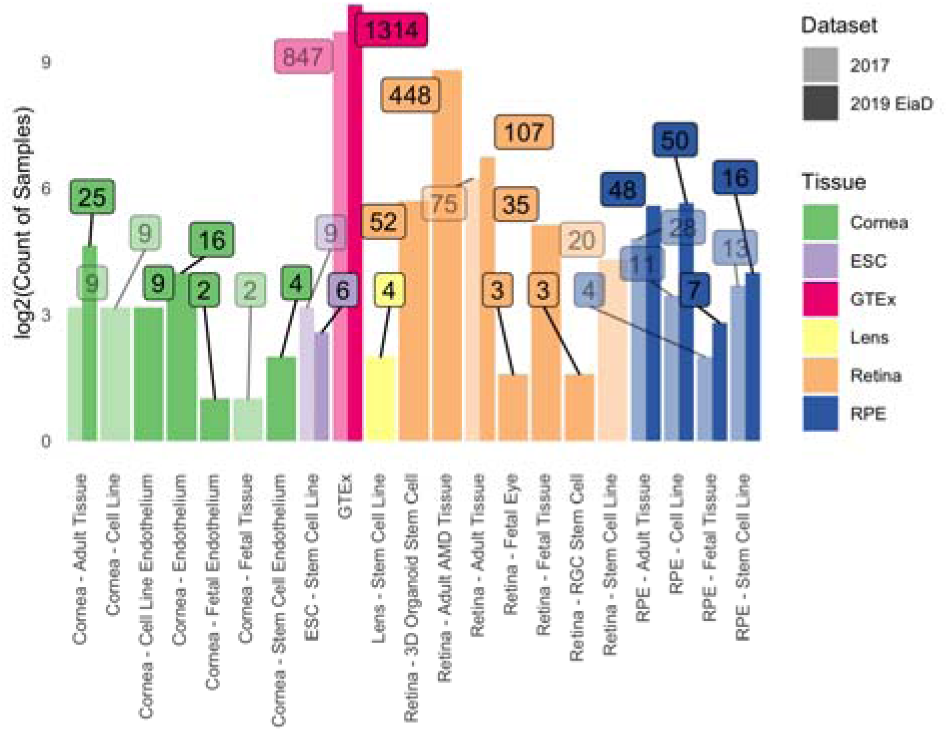
Substantial increase in eye tissue count and type from 2017 (180, lighter color) to 2019 (835, darker color) EiaD. We also improved the metadata labelling, the cornea samples (green) now delineates endothelial and epithelial tissues and the retina samples (orange) distinguish retina organoid and retinal ganglion cell (RGC) from stem cells. Counts for each bar plot given in the boxes. The y-axis is a log2 transformed count of samples passing our QC filters.

**Table 1:** EiaD contains a large set of diverse eye tissues, including embryonic stem cells (ESC). Eye lid and retina endothelium samples were included, but all failed to pass our QC filters.

| Tissue | Pre QC Count | Count | Sub-Tissue Types (Count) |
| --- | --- | --- | --- |
| Cornea | 62 | 56 | Adult Tissue (25), Cell Line Endothelium (9), Endothelium (16), Fetal Endothelium (2), Stem Cell Endothelium (4) |
| ESC | 12 | 6 | Stem Cell Line (6) |
| Eye lid | 4 | 0 |  |
| Lens | 9 | 4 | Stem Cell Line (4) |
| Retina | 681 | 648 | 3D Organoid Stem Cell (52), Adult Tissue (107), Adult Tissue AMD MGS 2 (172), Adult Tissue AMD MGS 3 (112), Adult Tissue AMD MGS 4 (61), Adult Tissue MGS 1 (103), Fetal Eye (3), Fetal Tissue (35), RGC Stem Cell (3) |
| Retinal Endothelium | 4 | 0 |  |
| RPE | 144 | 121 | Adult Tissue (48), Cell Line (50), Fetal Tissue (7), Stem Cell Line (16) |

Stem cell-derived cornea, stem cell-derived lens, and fetal retina are three new types of sub-tissues that are now available in EiaD. We have also substantially improved the granularity of the cornea tissue metadata, now delineating whether the tissue is from the endothelium or epithelium (Figure 1); previously these had been grouped together as adult tissue. Another substantial addition to the 2019 EiaD are non-protein coding genes; while protein-coding is the most common gene and transcript typse, there are dozens of different non-coding classes. The 2017 version of eyeIntegration only quantified protein coding genes and transcripts. We now quantify expression across 41 gene and 45 transcript types, including protein coding, retained intron, lincRNA, antisense, and pseuodogenes (Supplemental Table 2).

We have also added the large retina AMD post-morten Ratnapriya et al. cohort to EiaD 2019^37^. This cohort contains hundreds of samples ranging from non-AMD (Minnesota Grading System (MGS) 1) to severe AMD (MGS 4). While eyeIntegration is intended to be a source for normal tissues, we have made an exception for this study, as this is a large cohort and AMD is a common disease. We found our corrections methods did not group the non-AMD Ratnapriya et al. samples with our other collected retina samples (see Retina MGS in Supplemental Figure 3). This may be related to the lower mapping rate of the Ratnapriya et al. data (see Retina MGS in Supplemental Figure 2).

### 467 more GTEx samples and 9 new GTEx body sub-tissue types added to 2019 EiaD

Our previous dataset for eyeIntegration version 0.6 held about 20 samples per GTEx tissue type. We ran power calculations to assess our ability to detected >= 1 log2(Fold Change) in gene expression between two conditions to determine whether this is a sufficient number of samples (Supplemental Figure 4). Our calculations suggest, for example, that we have 83% power to detect a 1 log2(Fold Change) difference in gene expression with two groups of twenty samples. To increase our power to make significant eye to body comparisons, we added about 10 more samples per GTEx tissue types (which at 30 samples, would give about 90% power). We also took this opportunity to add bladder, bone marrow, cervix uteri, fallopian tube, ovary, prostrate, testis, uterus, and vagina GTEx tissue samples (Supplemental Table 1).

### Rigorous quality control and reproducible workflow system ensures high quality transcriptomes that consistently cluster together by tissue type

We built an automated pipeline for processing and analyzing all data for the web app using the program Snakemake, a python-based workflow management system that allows for efficient parallel execution of the analysis, facilitates reproduction by others, and simplifies long term maintenance of the EiaD data (Figure 2, Supplemental Figure 5)^57^. To create a high quality final dataset across the 2291 initial samples (Supplemental Table 3) and 67,315,523,736 reads we developed a rigorous quality control procedure as part of our analysis, considering a sample’s read mapping rate and median count level as well as behavior relative to samples of the same sub-tissue type (see Methods). To identify tube mislabeling or sample extraction issues, we used sample-level gene correlatio metrics (see methods) to identify variability within samples of the same sub-tissue and ensure overall consistency in data processing (Figure 3). After these steps 81 eye samples and 61 GTEx samples were removed.

**Figure 2:**
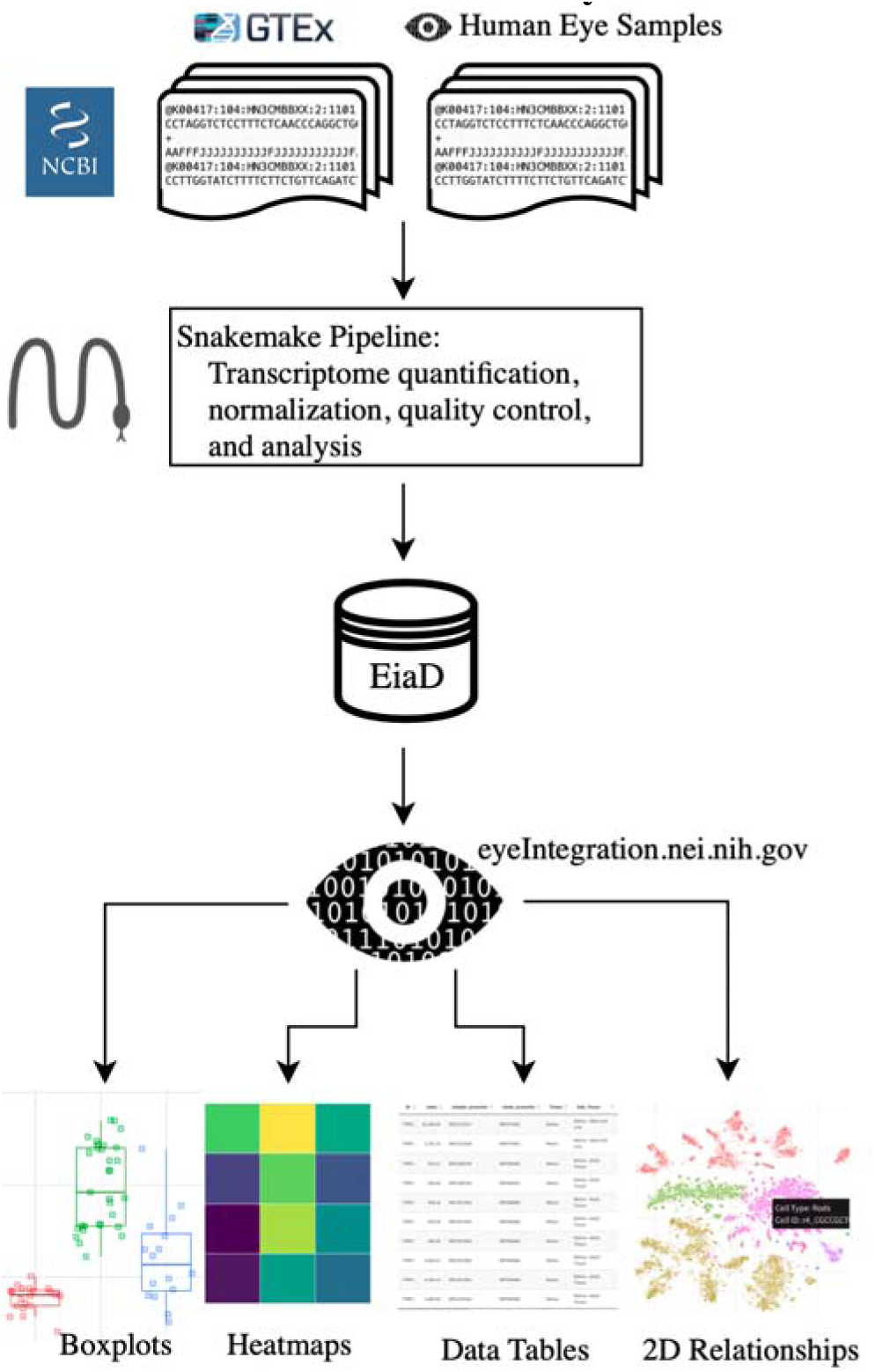
Raw RNA-seq data from the SRA is run through our pipeline to create the EiaD, which is used by eyeIntegration app to serve interactive gene expression visualizations across 73 tissues

**Figure 3:**
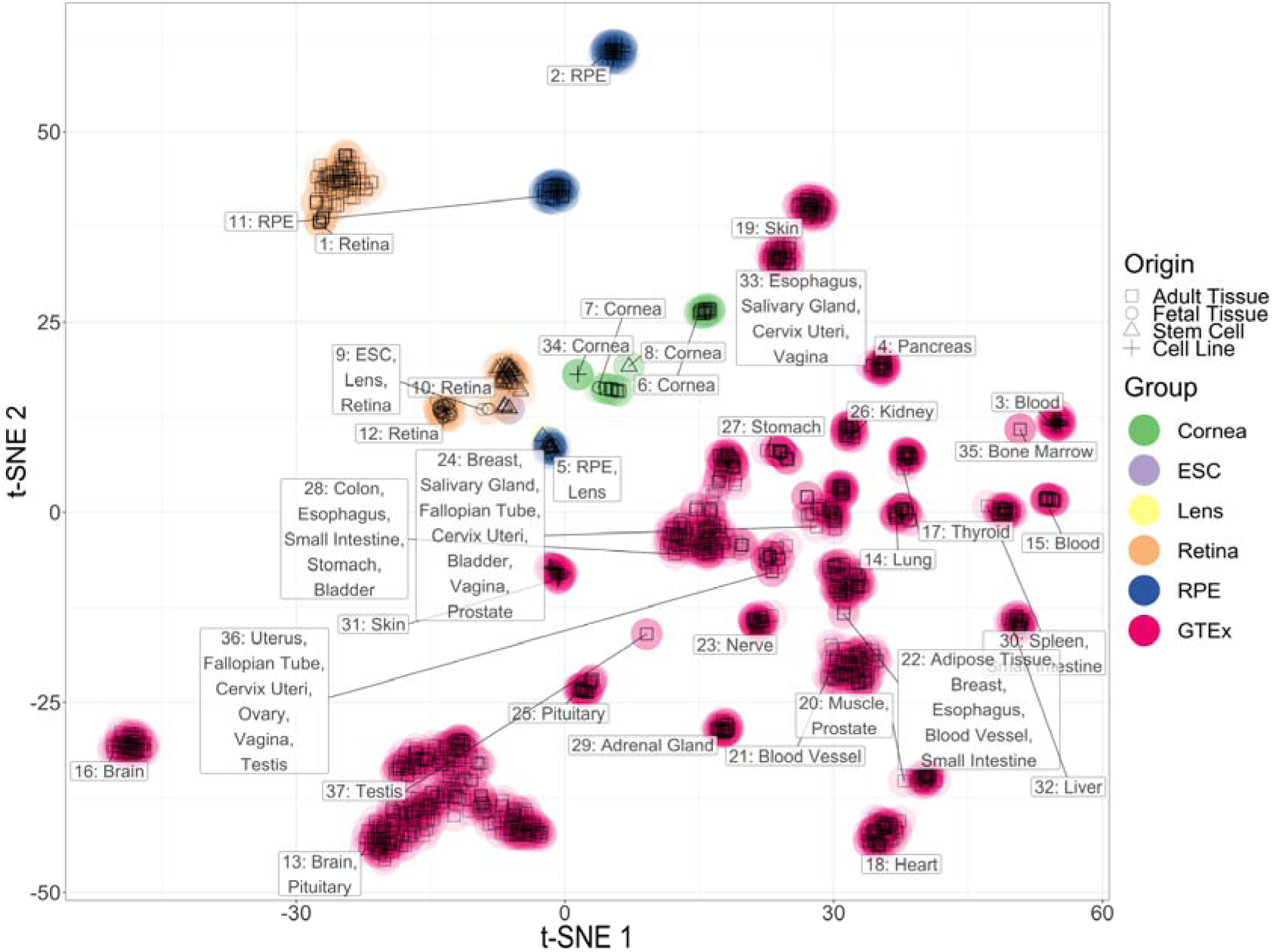
t-SNE two-dimensional transcriptome profiles by sample demonstrate effective quality control and transcriptome processing. Colors match different tissue types and shapes of points define the origin of the tissues.

To ensure there are no substantial differences in quantification of gene TPM values, we calculated the ^s^ between GTEX and EiaD generated TPM values for our shared GTEx samples (see methods); we computed an ^s^ of 0.89. Zhang et al. report that RNA-seq quantifications done between alignment-free methods (used in EiaD) and alignment-based methods (used by GTEx) get a ^s^ ranging from 0.89 to 0.93. As Zhang et al. compared quantification methods with identical gene references (we use Gencode GRCh38 gene models and GTEx uses hg19) and did not scale TPM score differently, our result falls in line with expectations.

After our quality control and processing workflow, we found that samples of the same tissue type and origin cluster well together (Figure 3 and Supplemental Table 4). For example, in the retina group, primary adult tissue clusters tightly and distinctly from other cell types, and retinal organoids and fetal retina samples cluster together. Our ability to uniformly cluster data by known biological source independent of study origin demonstrates that our workflow can effectively account for technical variation between studies.

While t-SNE is a powerful algorithm for grouping samples, it is not consistent for determining the relationships between clusters^61^; PCA is more useful in this regard. We ran a PCA dimensionality reduction (Supplemental Figure 6) on all samples, finding that the eye tissues still generally group together and apart from all other human body tissues. Adult retina is most similar to the brain tissue. RPE and cornea are most similar to blood, bone marrow, and skin.

### The eyeIntegration web app provides interactive visual portal to all data

The EiaD 2019 dataset is used directly by the eyeIntegration web app (https://eyeIntegration.nei.nih.gov). The web app was designed to provide a simple interface that has the same general concept – select specific genes and tissue and view relevant information. The web-app is divided into four general categories: expression, two-dimension sample relationships, gene networks, and data tables.

### Custom gene and tissue expression boxplots

The ‘Expression’ tab of the webpage provides a wealth of information about both gene- and transcript-level expression for eye and body tissues, giving the user the ability to compare the expression of different genes within a single tissue, as well as the expression of genes across multiple tissues (Figure 4A). The user first selects either the 2017 or 2019 gene or transcript EiaD dataset, then Hugo Gene Nomeclature Committee (HGNC) genes names (or ENSEMBL transcripts), then tissues. A boxplot is then generated after hitting the “Re(Draw) Plot” button with overlaid individual data points. On mouse-over, the metadata for the individual sample is displayed. A tabular report is generated based on selected genes and tissues: a table with links to Ensembl, GeneCard, and OMIM for each gene for quick referencing, and a table containing expression levels for each selected gene in each selected tissue. The tables can be arranged or sorted to the user’s preference and can be easily downloaded for local use.

**Figure 4:**
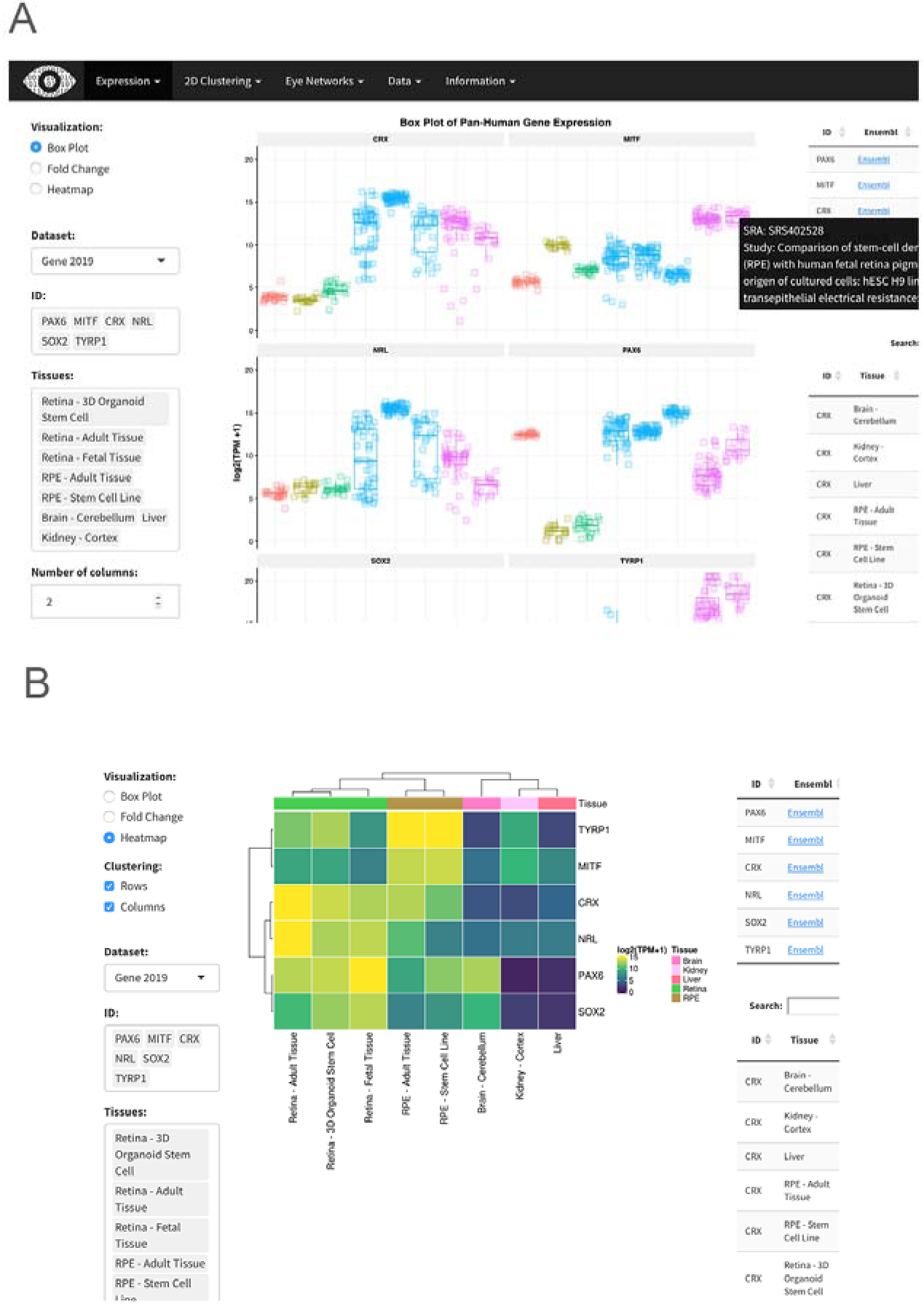
Screenshots from eyeIntegration web app. A. Pan-tissue gene expression box plots with accompanying data tables. The data tables display the rank (lower is more highly expressed) of each gene in each sub tissue, the decile of the rank (10 is the highest decile of expression), and the gene’s mean log2(TPM + 1) score for each sub tissue. B. Heatmap visualization.

Heatmap built by the R package ComplexHeatmaps based on expression can be drawn for selected genes and tissues and gene expression can be compared across many genes and tissues (Figure 4B)^62^. Finally, a session can be saved or shared by building a custom link for the session with the “Build URL Shortcut” button.

### Differential expression and gene ontology enrichment tests allow quick comparison of gene differences between groups

We performed multiple differential comparisons at the sub-tissue level within all eye tissues and against a pan-body synthetic set comprised of a stratified sample of all tissues present in our subset of the GTEx dataset, allowing quick identification of eye specific genes across 882 different comparisons. We have expanded the differential tests in the 2019 EiaD by adding the GTEx tissues as direct comparisons to our eye sub-tissues. The user can view the results selecting ‘Differential’ under the ‘Expression’ tab (Supplemental Figure 7D). As with ‘Expression’, the user can select which version of the web app to draw data from as well as select for gene- or transcript-level comparisons. The user additionally has the option to select different gene classes to examine, e.g. protein coding, lincRNA.

The results of differential expression are presented in a tabular format showing log_2_ fold change, average expression, and p-values. Depending on the comparison, there are 1 to 33380 differentially expressed genes (Supplementary Files). The table can be easily searched for any given gene, viewed and ordered to the user’s preference, and downloaded in CSV format. Differential expression can be visualized through fold change bar graphs with the ‘Pan-tissue plots’ selection under ‘Expression’. Additionally, we performed GO enrichment for all differential comparisons. Enriched GO terms are presented first as a word cloud, for quick comparison of GO enrichment. We provide tables, with similar viewing options as the differential expression table, for enriched GO terms in each class of a given differential comparison.

### Murine scRNA-seq enables testing of retina cell type specific expression

We incorporated scRNA-seq data from murine retina across two studies^58, 59^. This allows researchers to quickly examine gene expression across individual cell types in the retina. Single cell gene expression data is visualized through a heatmap showing the expression of a gene across multiple retinal cell types and different developmental time points, from embryonic day (E)11 to postnatal day (P)14 (when available), and a table of expression values is generated containing the expression data used to draw the heatmap (Figure 4C). We also provide t-SNE/UMAP based clustering using cell type specific labeling created by the publishing authors (Figure 4D, see Methods). The plots show all cell types present at a given developmental stage, and highlights cells expressing a gene above a user-selected given level.

### EiaD 2019 suggests that iPSC-derived organoids and fetal retina have closely related transcriptomes

There are, currently, two major approaches to studying developing human retina: post-mortem fetal tissue and stem-cell derived organoids. We looked at how well these approaches to studying developing retina compare at a transcriptomic level, both for tissue - organoid relationships and how well they correlate across early development.

To evaluate how the tissues and organoids compare at a transcriptome level, we looked at the same t-SNE plot from Figure 3 and focused in on the three types of retina tissue (adult, fetal, and organoid) (Figure 5A). Here we saw three distinct groupings: adult retina (1), developing fetal retina and stem cell-derived organoid (2), and undifferentiated and early differentiating stem cells (3). We identified several organoid samples in cluster 3, but these share one important difference from the rest of the organoid samples in cluster 3: they have been differentiating for less than 30 days (shape ‘X’). All of the organoid retina samples in cluster 2 are older than 50 days.

**Figure 5:**
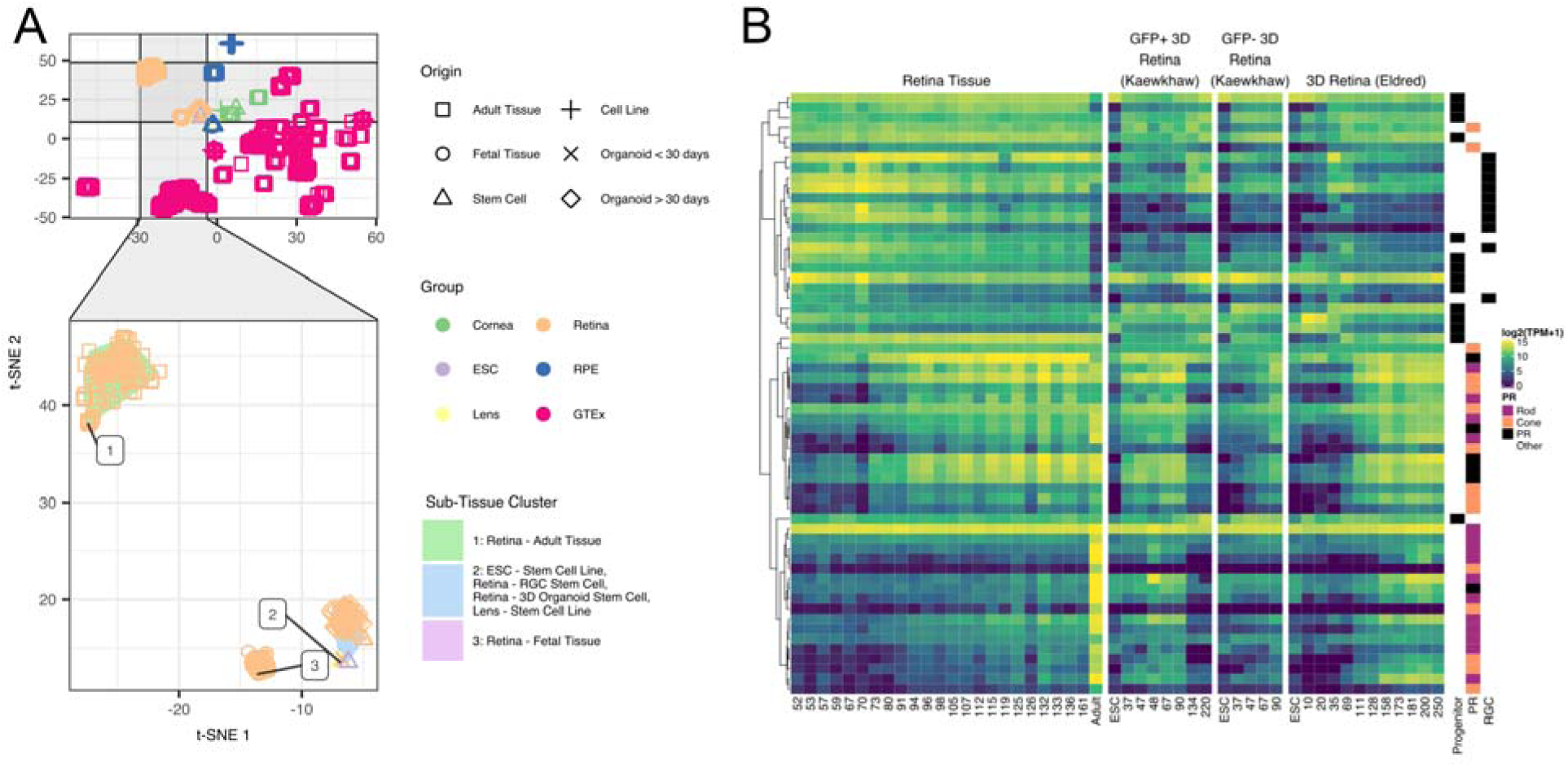
Organoid retina, stem cell retina, and fetal retina tissue have highly similar transcriptomes. The zoom inset (A) shows the retina samples. The “Sub-Tissue Cluster” shading shows the cluster membership of the three major groups. The shapes of the points show the different origin types - notable types include the square for adult, the ‘X’ for organoid under 30 days of differentiation, and the diamond for organoid over 30 days of differentiation. Major markers of retina progenitor, photoreceptors (cone and rod), and retinal ganglia cells (RGC) have similar gene expression patterns across development in retina fetal tissue and organoids.

To assess how similarly the fetal and organoid retina develop through time, we plotted expression of retinal progenitors, photoreceptors, and retinal ganglion markers by time in days (Figure 5B). Each row is a gene marker of either retinal progenitor, photoreceptor, or RGC. The rows are hierarchially clustered to put more similar expression patterns closer together, as denoted by the height of the dendrogram. We split the organoid tissues into three groups: Kaewkhaw et al. GFP+ and GFP-samples, and Eldred et al. samples^12, 63^. The Kaewkhaw samples are flow sorted for a GFP marker (GFP+) under the control of the *CRX* promoter, an important regulator of photoreceptor development. GFP+ cells wouldbe enriched in photoreceptor populations. We saw that the retinal progenitor, photoreceptor, and RGC groups are largely clustered together, with patterns of expression consistent across the fetal retinal and organoid groups.

### Differential gene expression of organoid retina versus fetal tissue identifies sets of genes relating to patterning (HOXB family), cell adhesion (protocadherin family), and RGC identity (BRN3/POU4F, NEFL, GAP43, SNCG)

To identify specific changes between retinal organoid and fetal retina tissue, we performed differential gene expression and GO term enrichment analyses. The GO term enrichment identified cell adhesion (protocadherins) and patterning (HOXB family) as enriched gene sets in retinal organoids As there is some evidence suggesting that protocadherins influence RGC viability and we noticed that several RGC markers appeared to have lower expression in the organoids compared to the fetal tissue Figure 5B we looked more closely into RGC marker expression^64^.

We plotted HOXB family, protocadherin family and RGC genes in a heatmap visualization, with columns as age in days of fetal or organoid retina. Rows are genes, split by the three different groups of genes and are internally clustered by how similar the expression patterns are. We observed that there are strong, consistent gene expression differences in these three groups of genes between fetal retina and the organoid samples (Supplemental Figure 8). We also plotted the differential expression values between all organoids and all fetal retina samples; all genes across all three groups are significantly differentially expressed with an FDR corrected p value < 0.01.

### Limitations of the RNA-seq quantification in eyeIntegration

Salmon quantification, while highly performant and accurate, has a higher variance for lower read depth samples and shorter transcripts^54^. Extra care should be taken with comparisons with lower counts of samples (cornea, RGC) as smaller sample numbers decrease the confidence in differential expression. We do not recommend you directly compare our TPM values with your counts data as there are many important variables that will differ. Instead run our Snakemake pipeline (https://www.github.com/davemcg/EiaD_build), adding your samples. Finally, we would like to remind any users that RNA-seq methods measure mRNA levels, but the functional unit is the protein; westerns are still the gold standard with which to evaluate expression and localization.

### Data accessibility

Individual data files for gene expression and sample metadata can be downloaded from the ‘Data’ tab on the web app. All data and code used to generate the web app can be installed from the R command line by running devtools::install_github(‘davidmcg/eyeIntegration_app’). The code for the EiaD data processing pipeline can be found at https://github.com/davemcg/EiaD_build.

## Discussion

EiaD 2019 contains a large set of carefully curated, reproducibly processed human eye RNA-seq datasets alongside a human body tissue comparison set from the GTEx project. It is available for local install as an R package at https://www.github.com/davemcg/eyeIntegration_app and it is served via a web app, eyeIntegration at https://eyeIntegration.nei.nih.gov. The web app offers a wide range of user-driven visualizations to compare expression of genes across dozens of human body and eye tissues. Furthermore, murine scRNA-seq datasets have been incorporated, allowing for examination of retina cell type-specific gene expression. Several human and non-human primate studies have been posted in the past year on the pre-print server bioRxiv and as the raw data becomes publicly available, we will be updating this section of eyeIntegration^65–67^.

If you wish to have your data added to EiaD in the future, we suggest you 1. deposit data into GEO/SRA, 2. use clear, descriptive, consistent, and detailed metadata for each sample, and 3. (optional) contact the corresponding author. Contacting the corresponding author is only necessary if you feel your data should be included in EiaD and was deposited into the SRA before May 8th, 2019.

As human fetal tissue is difficult to obtain and thus not very amenable for chemical or genetic modification, it is crucial for organoid-based models to be developed. Our merging of these datasets and analysis at the transcriptome level (as compared to cross-analyzing using a limited number of known marker genes) indicates that these two approaches successfully recapitulate fetal retina tissue, to a first approximation, at the whole transcriptome level. However as organoids do not develop to full function, it is important to look at how gene expression differs between retinal organoid and fetal tissue so as to suggest areas for improvement.

We used our large dataset to narrow in on three core processes which differ significantly and substantially between retinal organoids and fetal retina. First we showed that the HOXB family is overexpressed in the organoids. The homeobox family is well known to initiate polarity of the embryo during early development^68^. Retinoic acid is applied at about day 20 in culture to help differentiate stem cell to organoids and is also known to activate genes members of the HOXB family. The lack of HOXB expression at any age in fetal retina and the broad chromatin and gene expression changes HOXB family members can mediate suggests that HOXB activity may be unwanted for organoid maturation.

Next, we detected several protocadherins more highly expressed in the fetal tissue, relative to the organoids. Protocadherins mediate cell to cell connections and, in the developing mouse, are shown to be important for spinal internneurons and RGC survival^64, 69^. We would predict that decreased protocadherin expression reduces the number and maturation of RGC. Indeed we observed that many canonical RGC markers, while present in detectable levels in the organoids, are signficantly underexpressed relative to fetal tissue. This result suggests that modifying culture conditions to promote protocadherin expression may result in higher RGC yield and survival.

We built the Eye in a Disk dataset and the accompanying web app, eyeIntegration in the hopes that easily accessible gene expression across tissue space and time will be a useful tool for hypothesis generation and refinement in eye research. Wrapping all of the data processing steps in a Snakemake pipeline has several important advantages for the community: our code is publicly available for review, our analyses are reproducible, future sample updates can be streamlined in with less effort, and because all the processing is in modular pieces it is easier to add new analysis steps. In the future, we plan on regularly adding new samples to EiaD, offering *de novo* eye tissue transcriptomes, expanding the single cell RNA-seq expression tooling, adding non-human eye samples, and epigenetic datasets.

## Supporting information

Supplemental File 2.xls

Supplemental File 1.xls

power_calc.R

## Supplementary Figures and Tables

**Supplemental Figure 1:**
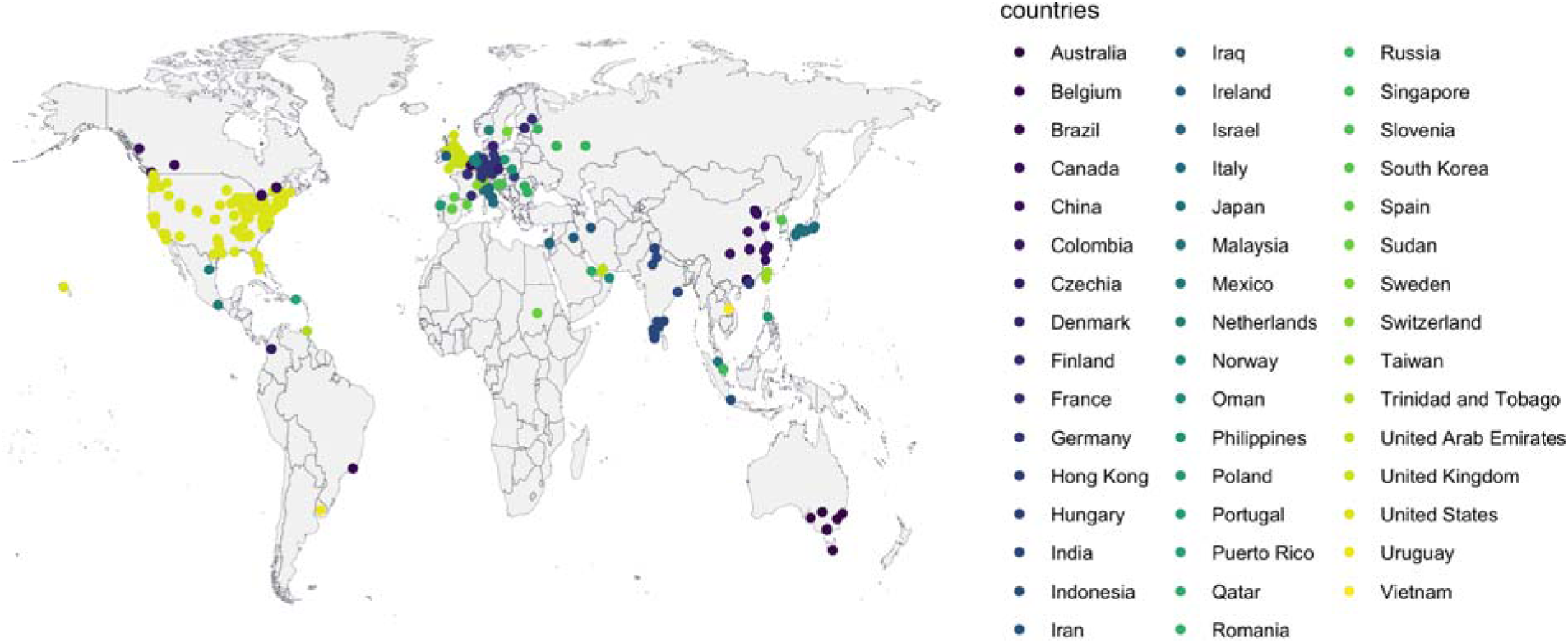
As of June 2019, eyeIntegration has had usage across 367 cities and 47 countries.

**Supplemental Figure 2:**
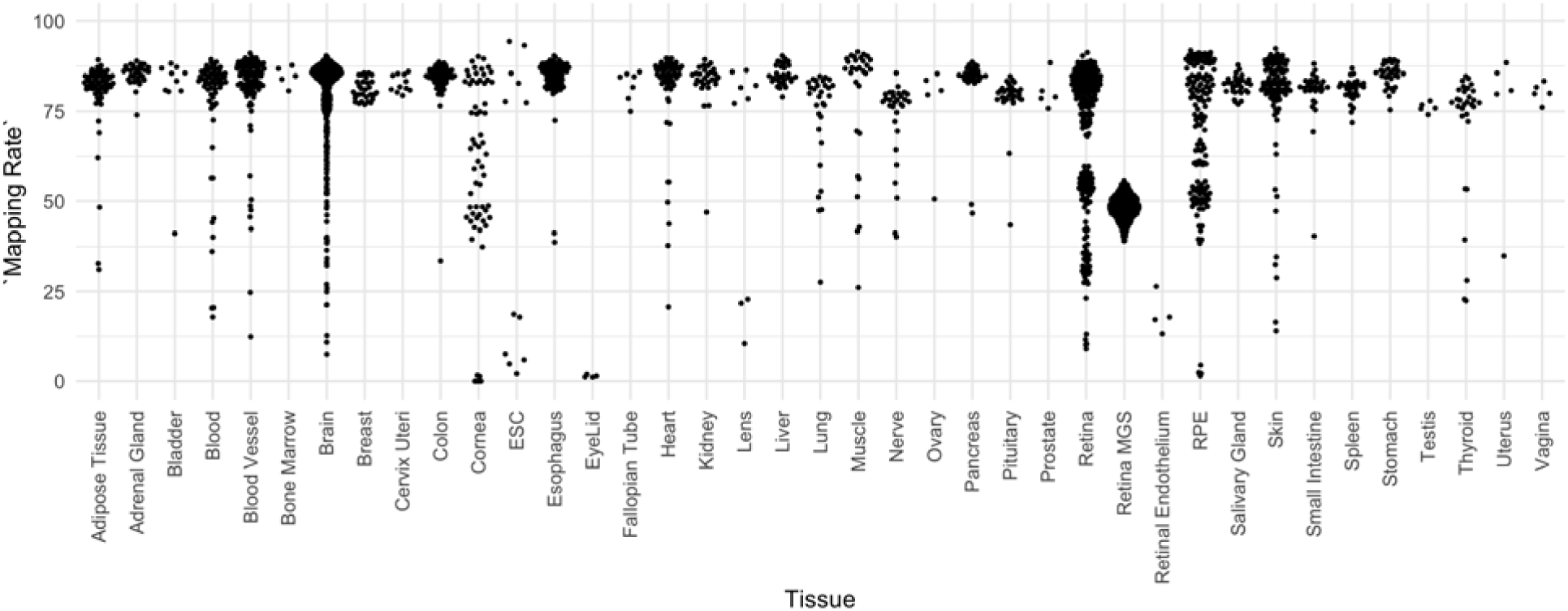
Salmon mapping rate for each sample, grouped by tissue type. 1st quartile mapping rate is 51.5%, median is 80.5%, mean is 70.5%, and third quartile is 85.1%.

**Supplemental Figure 3:**
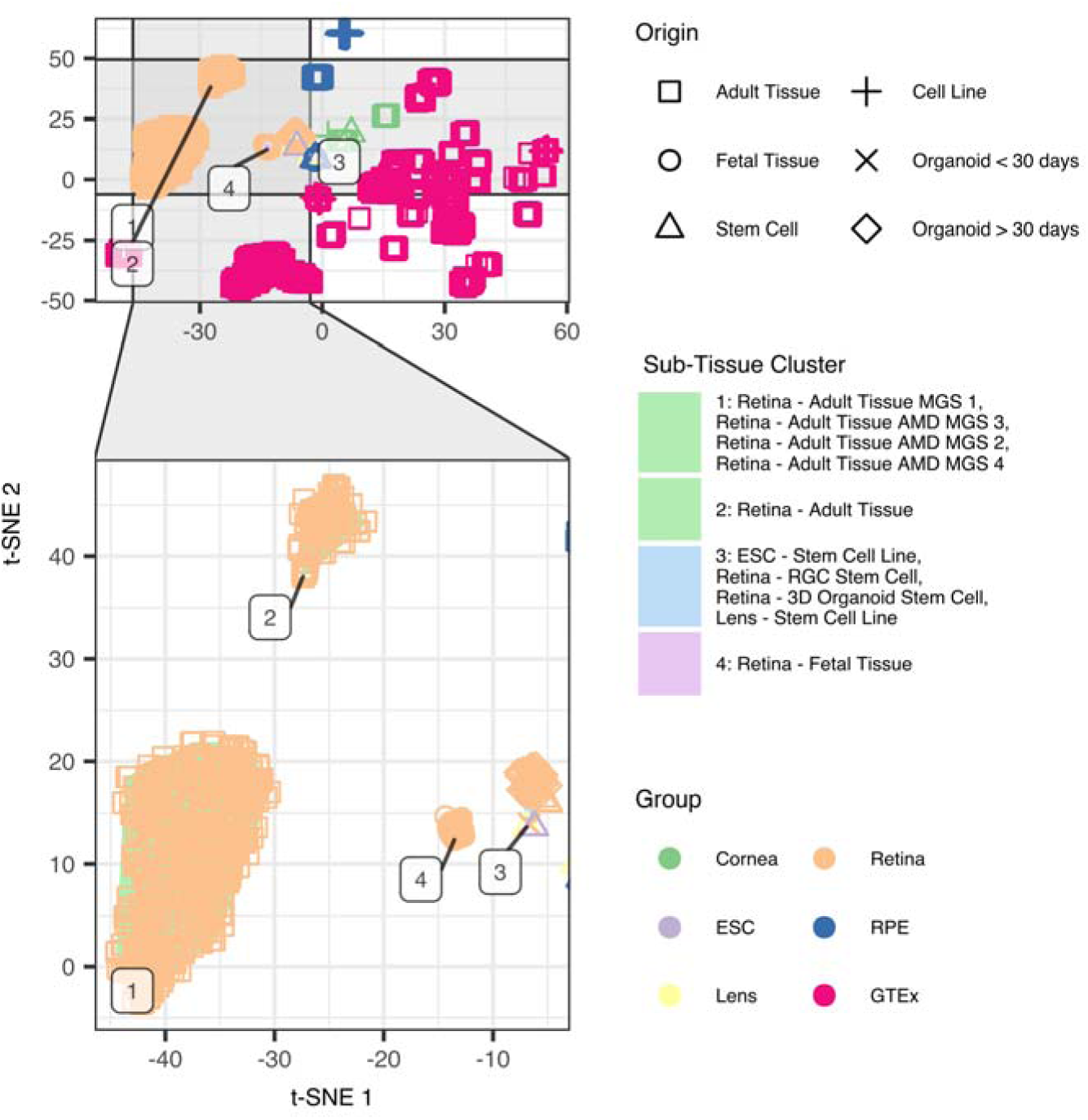
Adult retina samples from Ratnapriya et al. (MGS 1 - 4, not-AMD is MGS 1, AMD is MGS 2 through 4) cluster independently from all other adult retina samples collected.

**Supplemental Figure 4:**
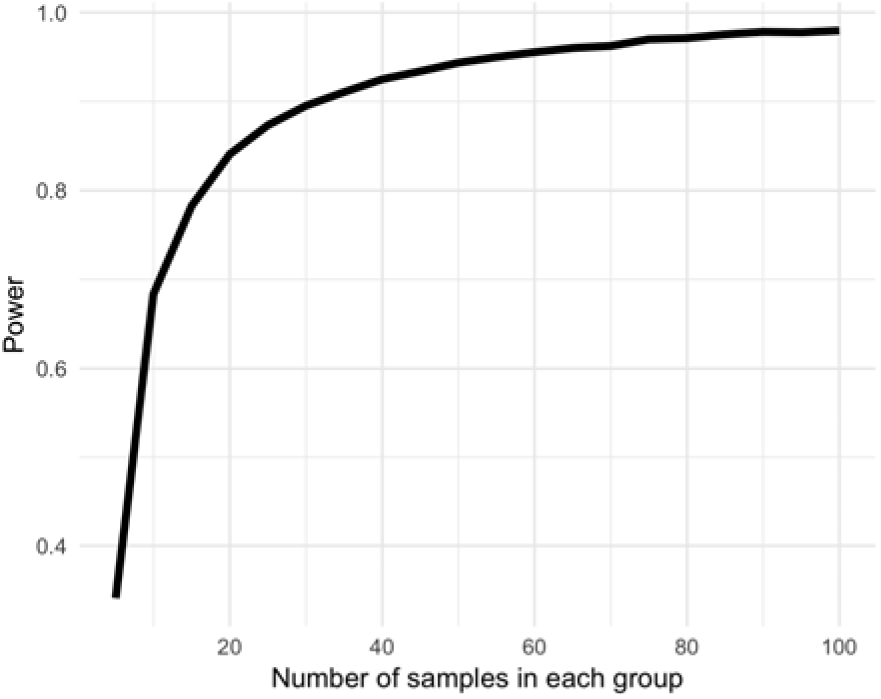
Power curve to assess ability to detect >= 1 log2(FoldChange) in gene expression between n samples (x-axis) in each group.

**Supplemental Figure 5:**
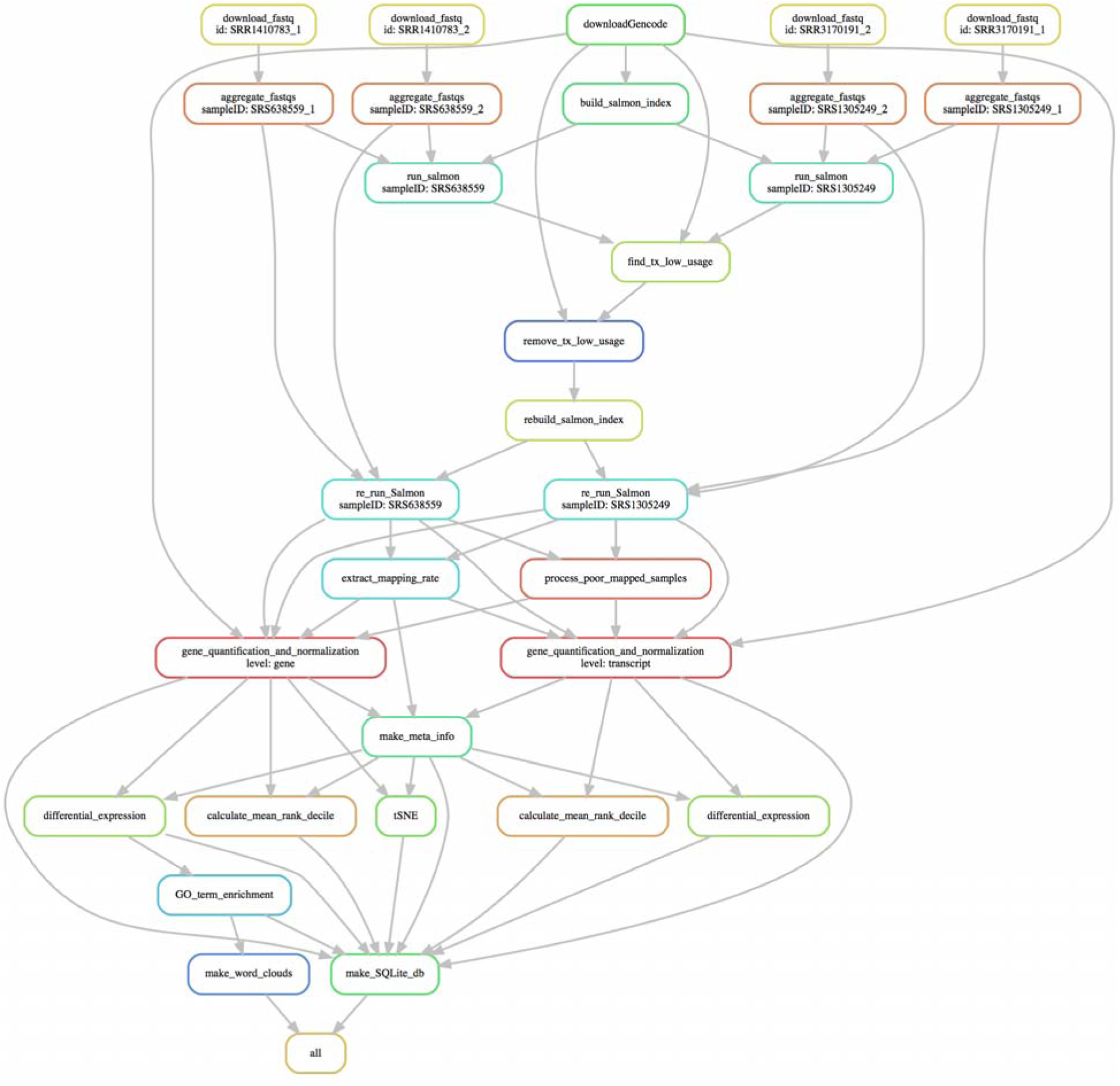
Snakemake pipeline to create EiaD 2019 consists of small modular compute sections to ensure sample tracking through the full pipeline

**Supplemental Figure 6:**
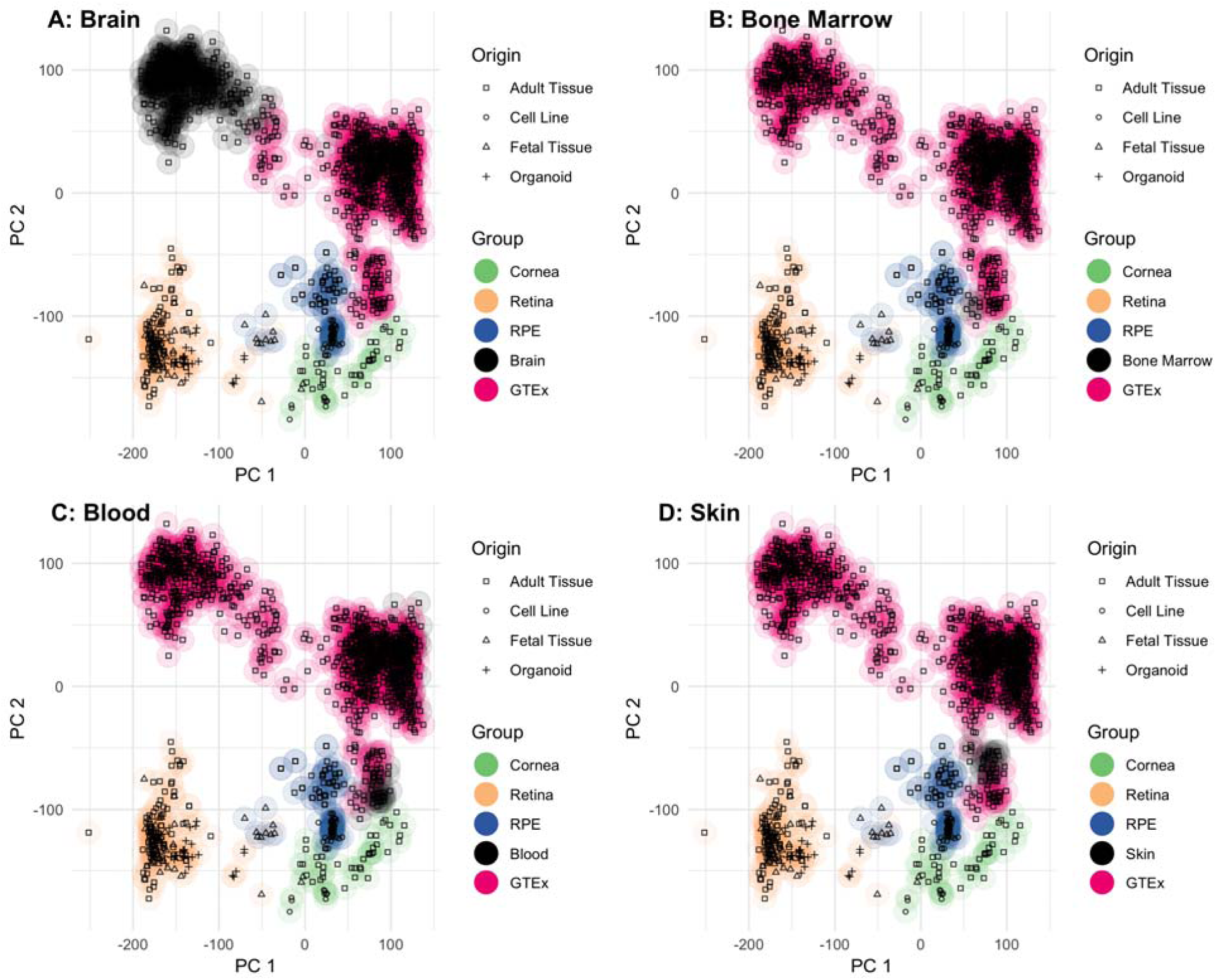
PCA plot of all samples suggests that the non-eye tissue most similar to adult retina is the brain, RPE and cornea are most similar to bone marrow, blood, and skin

**Supplemental Figure 7:**
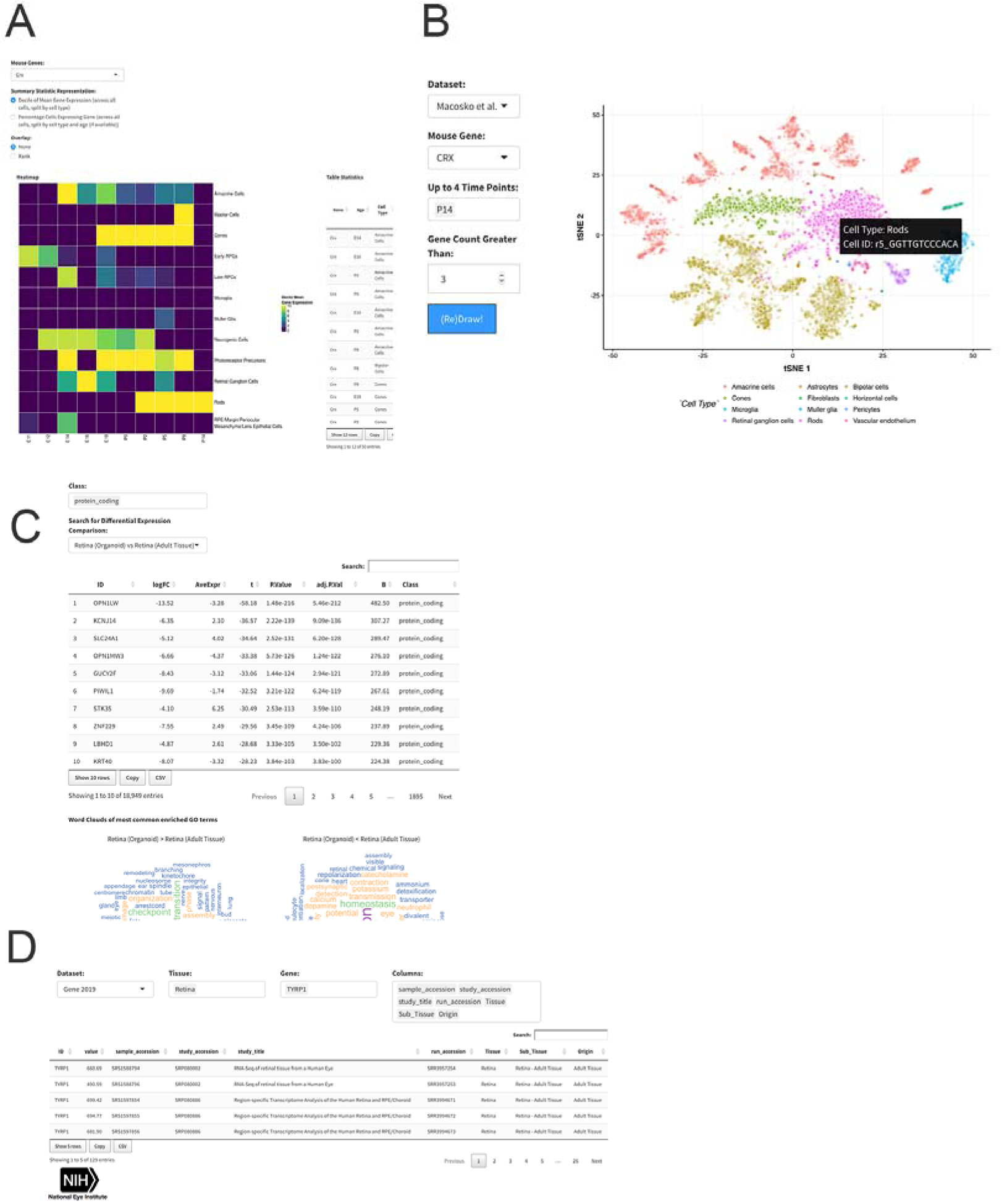
A. CRX mouse retina gene expression heatmap and table information from Clark et al. E11 to P14 scRNA-seq. B. t-SNE visualization of gene expression profiles of individual cells from Macosko et al. C. Data table export view. D. Differential gene expression across different tissues and with GO term enrichment.

**Supplemental Figure 8:**
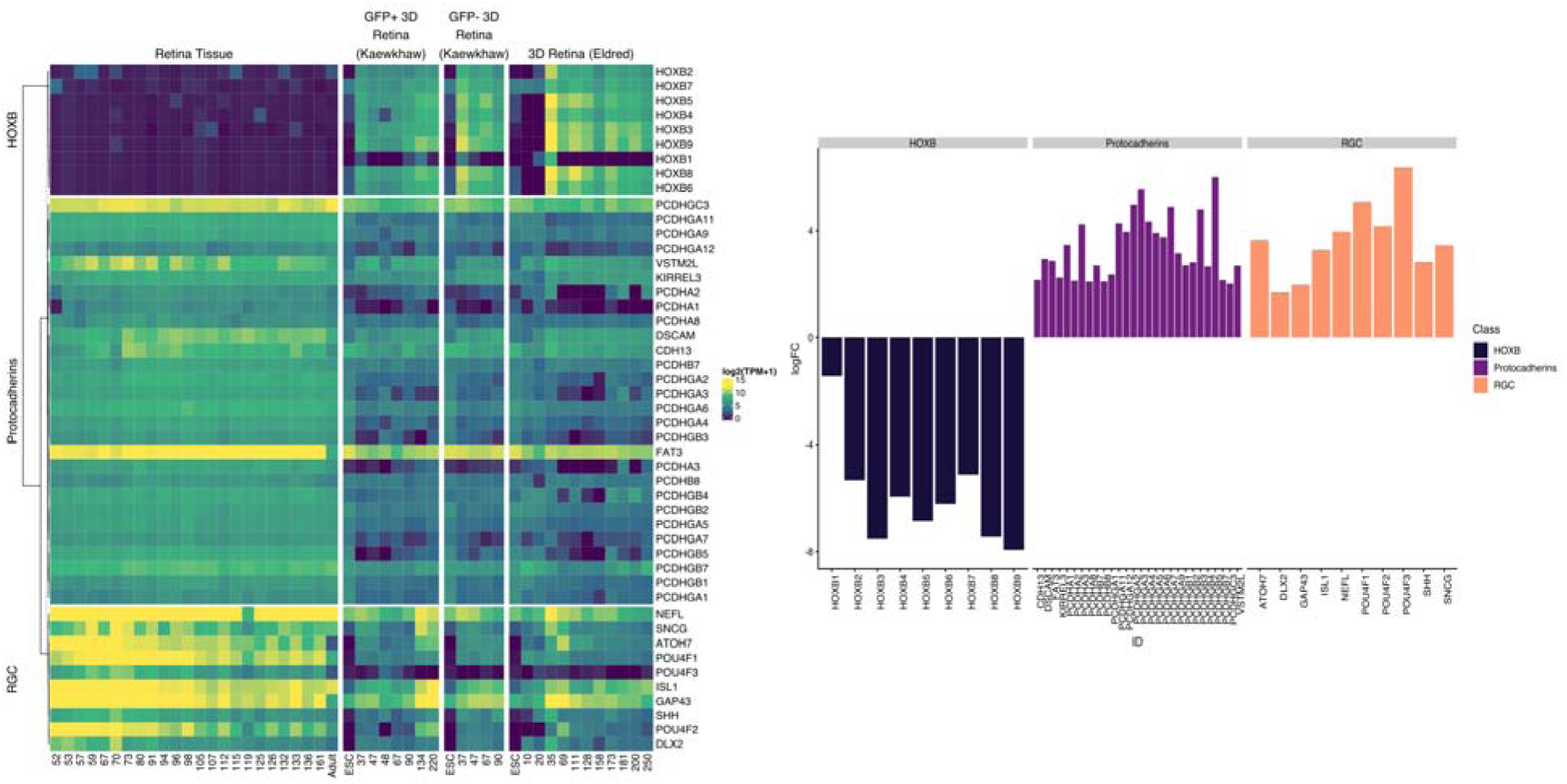
Heatmap of three sets of genes across fetal retina and organoids divided by age in days (A). Bar plot of differential expression from retina to organoid, where positive values are genes that are higher expressed in fetal tissue than organoid (B). All logFC expression values have FDR corrected p value < 0.01.

**Supplemental Table 1:** EiaD holds hundreds of GTEx tissues to provide a broad comparison set

| Group | Version | Pre QC Count | Count | Tissue Types (Count) |
| --- | --- | --- | --- | --- |
| GTEx | 2017 | 847 | 847 | Adipose Tissue (39), Adrenal Gland (20), Blood (26), Blood Vessel (57), Brain (255), Breast (20), Colon (40), Esophagus (59), Heart (37), Kidney (17), Liver (20), Lung (20), Muscle (19), Nerve (20), Pancreas (20), Pituitary (20), Salivary Gland (20), Skin (59), Small Intestine (20), Spleen (20), Stomach (19), Thyroid (20) |
| GTEx | 2019 | 1375 | 1314 | Adipose Tissue (54), Adrenal Gland (30), Bladder (9), Blood (49), Blood Vessel (88), Bone Marrow (5), Brain (372), Breast (30), Cervix Uteri (11), Colon (58), Esophagus (87), Fallopian Tube (7), Heart (58), Kidney (27), Liver (30), Lung (29), Muscle (29), Nerve (30), Ovary (4), Pancreas (30), Pituitary (29), Prostate (5), Salivary Gland (30), Skin (83), Small Intestine (29), Spleen (30), Stomach (30), Testis (5), Thyroid (27), Uterus (4), Vagina (5) |

**Supplemental Table 2:** Dozens of different types of gene and transcript types quantified

| Biotype | Gene Count | Transcript Count | Definition |
| --- | --- | --- | --- |
| bidirectional_promoter_lncRNA | 68 | 122 | A non-coding locus that originates from within the promoter region of a protein-coding gene, with transcription proceeding in the opposite direction on the other strand. |
| unitary_pseudogene | 22 | 21 | A species-specific unprocessed pseudogene without a parent gene, as it has an active orthologue in another species. |
| retained_intron |  | 6552 | Alternatively spliced transcript believed to contain intronic sequence relative to other, coding, variants. |
| protein_coding | 19012 | 34969 | Contains an open reading frame (ORF). |
| processed_transcript | 533 | 4635 | Doesn't contain an ORF. |
| antisense | 4014 | 5656 | Has transcripts that overlap the genomic span (i.e. exon or introns) of a protein-coding locus on the opposite strand. |
| pseudogene | 11 | 19 | Have homology to proteins but generally suffer from a disrupted coding sequence and an active homologous gene can be found at another locus. Sometimes these entries have an intact coding sequence or an open but truncated ORF, in which case there is other evidence used (for example genomic polyA stretches at the 3' end) to classify them as a pseudogene. Can be further classified as one of the following. |
| nonsense_mediated_decay |  | 3402 | If the coding sequence (following the appropriate reference) of a transcript finishes >50bp from a downstream splice site then it is tagged as NMD. If the variant does not cover the full reference coding sequence then it is annotated as NMD if NMD is unavoidable i.e. no matter what the exon structure of the missing portion is the transcript will be subject to NMD. |
| IG_C_gene | 14 | 17 | Immunoglobulin (Ig) variable chain and T-cell receptor (TcR) genes imported or annotated according to the IMGT. |
| IG_J_gene | 2 | 2 | Immunoglobulin (Ig) variable chain and T-cell receptor (TcR) genes imported or annotated according to the IMGT. |
| IG_V_gene | 127 | 127 | Immunoglobulin (Ig) variable chain and T-cell receptor (TcR) genes imported or annotated according to the IMGT. |
| TR_C_gene | 6 | 6 | Immunoglobulin (Ig) variable chain and T-cell receptor (TcR) genes imported or annotated according to the IMGT. |
| TR_J_gene | 6 | 6 | Immunoglobulin (Ig) variable chain and T-cell receptor (TcR) genes imported or annotated according to the IMGT. |
| TR_V_gene | 41 | 36 | Immunoglobulin (Ig) variable chain and T-cell receptor (TcR) genes imported or annotated according to the IMGT. |
| IG_C_pseudogene | 3 | 3 | Inactivated immunoglobulin gene. |
| IG_J_pseudogene | 1 | 1 | Inactivated immunoglobulin gene. |
| IG_V_pseudogene | 21 | 18 | Inactivated immunoglobulin gene. |
| TR_V_pseudogene | 4 | 4 | Inactivated immunoglobulin gene. |
| sense_intronic | 659 | 664 | Long non-coding transcript in introns of a coding gene that does not overlap any exons. |
| sense_overlapping | 153 | 180 | Long non-coding transcript that contains a coding gene in its intron on the same strand. |
| lincRNA | 4323 | 5881 | Long, intervening noncoding (linc) RNA that can be found in evolutionarily conserved, intergenic regions. |
| rRNA_pseudogene | 24 | 24 | Non-coding RNA predicted to be pseudogene by the Ensembl pipeline |
| miRNA | 130 | 130 | Non-coding RNA predicted using sequences from Rfam and miRBase |
| misc_RNA | 260 | 260 | Non-coding RNA predicted using sequences from Rfam and miRBase |
| Mt_rRNA | 2 | 2 | Non-coding RNA predicted using sequences from Rfam and miRBase |
| Mt_tRNA | 22 | 22 | Non-coding RNA predicted using sequences from Rfam and miRBase |
| ribozyme | 2 | 2 | Non-coding RNA predicted using sequences from Rfam and miRBase |
| rRNA | 5 | 5 | Non-coding RNA predicted using sequences from Rfam and miRBase |
| scaRNA | 15 | 15 | Non-coding RNA predicted using sequences from Rfam and miRBase |
| scRNA | 1 | 1 | Non-coding RNA predicted using sequences from Rfam and miRBase |
| snoRNA | 238 | 242 | Non-coding RNA predicted using sequences from Rfam and miRBase |
| snRNA | 104 | 106 | Non-coding RNA predicted using sequences from Rfam and miRBase |
| polymorphic_pseudogene | 19 | 22 | Pseudogene owing to a SNP/DIP but in other individuals/haplotypes/strains the gene is translated. |
| unprocessed_pseudogene | 593 | 585 | Pseudogene that can contain introns since produced by gene duplication. |
| translated_processed_pseudogene | 2 | 2 | Pseudogene that has mass spec data suggesting that it is also translated. |
| processed_pseudogene | 2537 | 2435 | Pseudogene that lack introns and is thought to arise from reverse transcription of mRNA followed by reinsertion of DNA into the genome. |
| transcribed_processed_pseudogene | 338 | 169 | Pseudogene where protein homology or genomic structure indicates a pseudogene, but the presence of locus-specific transcripts indicates expression. |
| transcribed_unitary_pseudogene | 101 | 38 | Pseudogene where protein homology or genomic structure indicates a pseudogene, but the presence of locus-specific transcripts indicates expression. |
| transcribed_unprocessed_pseudogene | 680 | 327 | Pseudogene where protein homology or genomic structure indicates a pseudogene, but the presence of locus-specific transcripts indicates expression. |
| TEC | 755 | 803 | To be Experimentally Confirmed. This is used for non-spliced EST clusters that have polyA features. This category has been specifically created for the ENCODE project to highlight regions that could indicate the presence of protein coding genes that require experimental validation, either by 5' RACE or RT-PCR to extend the transcripts, or by confirming expression of the putatively-encoded peptide with specific antibodies. |
| non_stop_decay |  | 10 | Transcript that has polyA features (including signal) without a prior stop codon in the CDS, i.e. a non-genomic polyA tail attached directly to the CDS without 3' UTR. These transcripts are subject to degradation. |
| 3prime_overlapping_ncRNA | 25 | 28 | Transcript where ditag and/or published experimental data strongly supports the existence of short non-coding transcripts transcribed from the 3'UTR. |
| non_coding | 2 | 2 | Transcript which is known from the literature to not be protein coding. |
| macro_lncRNA | 1 | 1 | Unspliced lncRNA that is several kb in size. |

**Supplemental Table 3:**
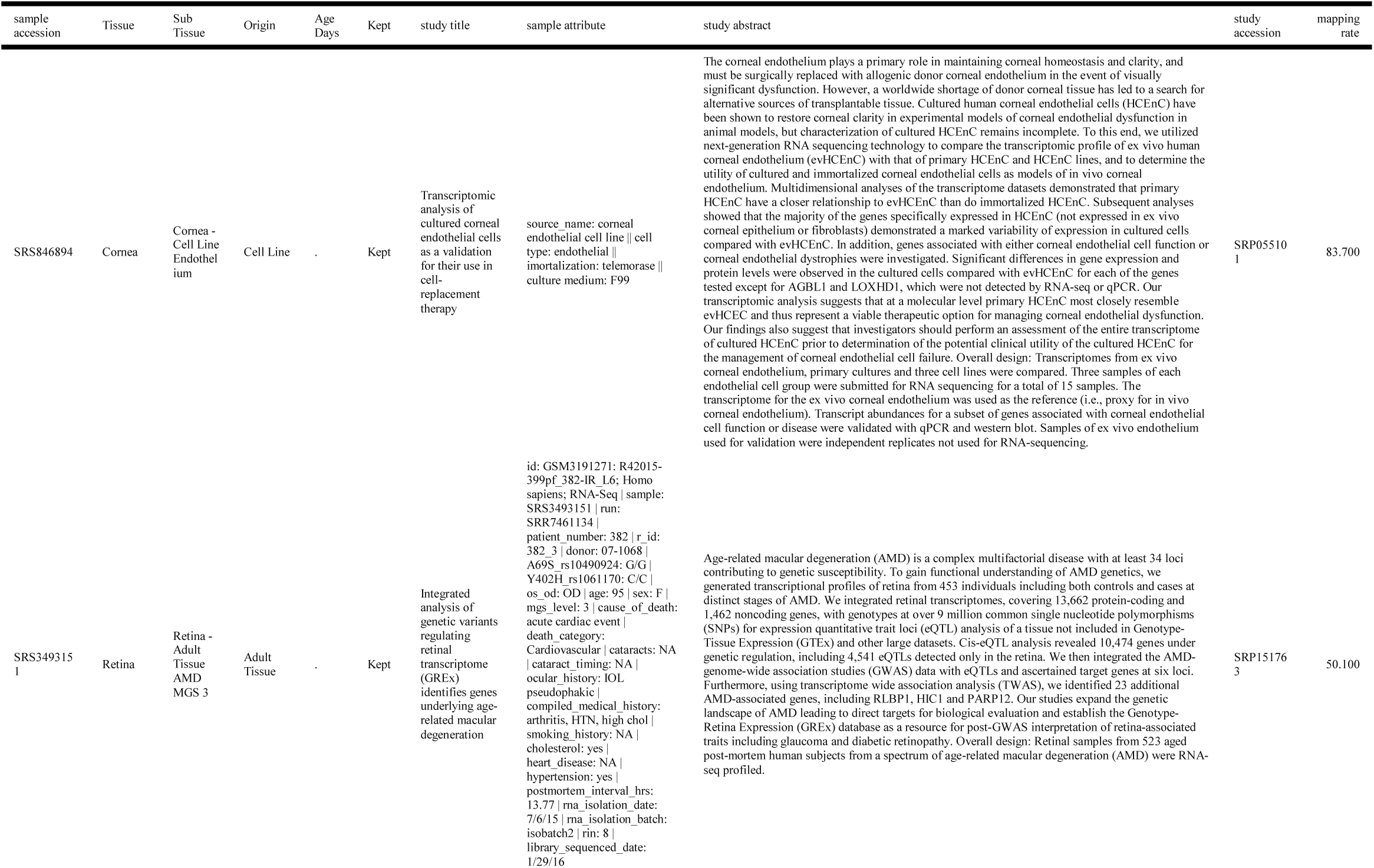
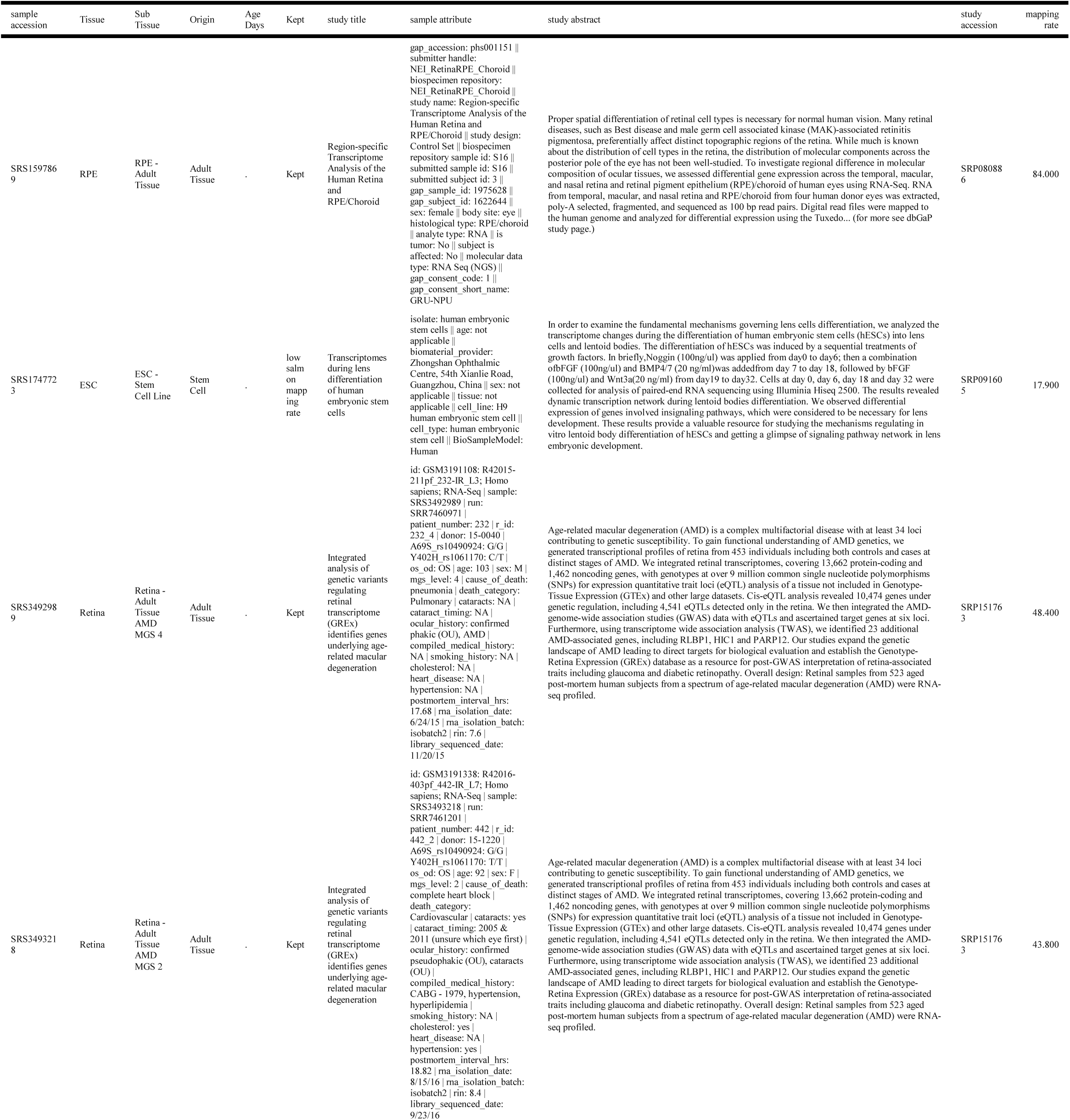
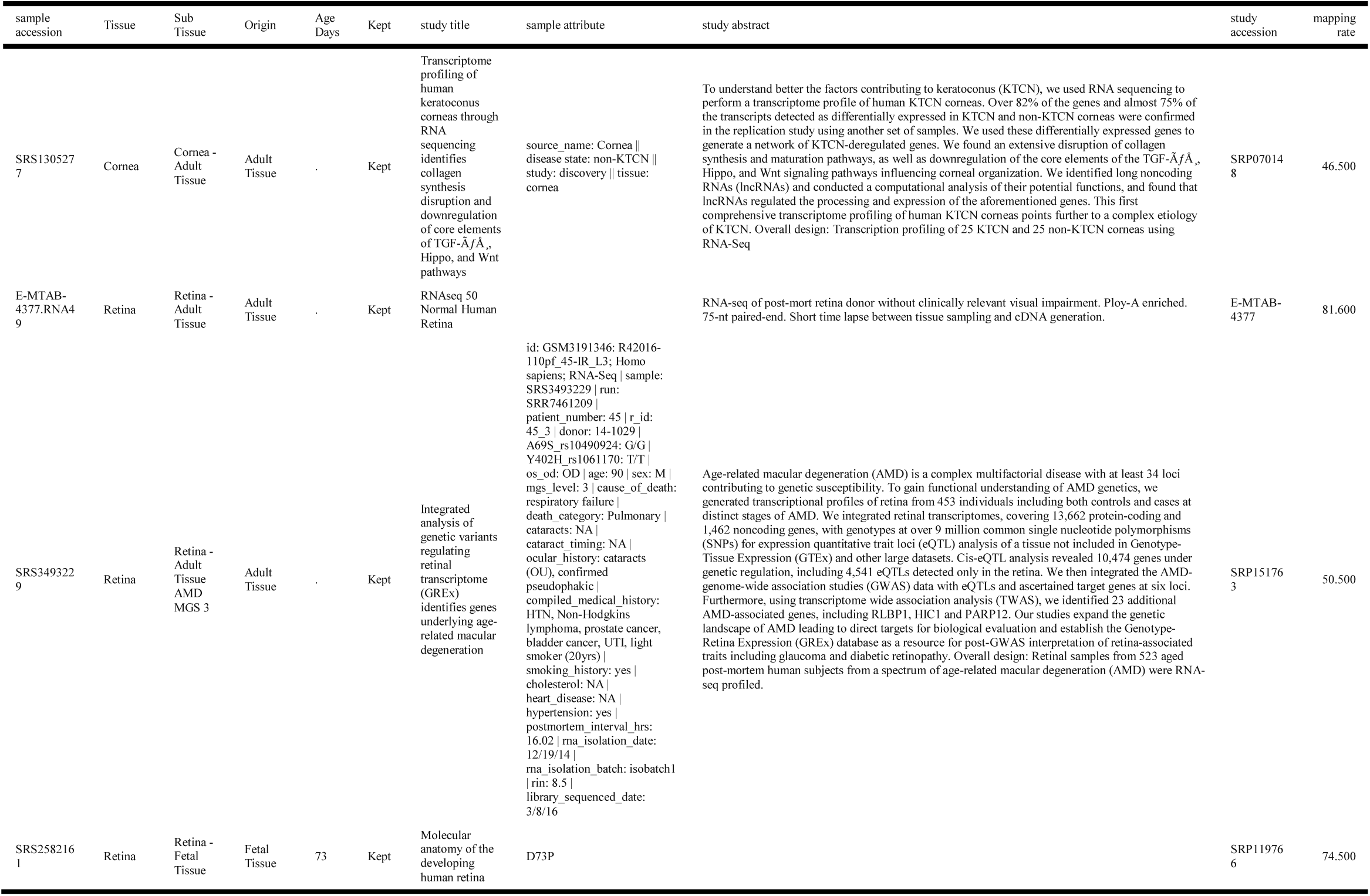
Full metadata for 10 random eye samples. Full metadata available as supplementary file “metadata.csv.”

**Supplemental Table 4:** Counts of Sub Tissues in each tSNE - dbscan based cluster group

| Sub Tissue | Cluster | Count |
| --- | --- | --- |
| Retina - Adult Tissue | 1 | 107 |
| RPE - Cell Line | 2 | 50 |
| Cells - EBV-transformed lymphocytes | 3 | 30 |
| Pancreas | 4 | 30 |
| Lens - Stem Cell Line | 5 | 2 |
| RPE - Fetal Tissue | 5 | 7 |
| RPE - Stem Cell Line | 5 | 16 |
| Cornea - Adult Tissue | 6 | 25 |
| Cornea - Endothelium | 7 | 16 |
| Cornea - Fetal Endothelium | 7 | 2 |
| Cornea - Stem Cell Endothelium | 8 | 4 |
| ESC - Stem Cell Line | 9 | 6 |
| Lens - Stem Cell Line | 9 | 2 |
| Retina - 3D Organoid Stem Cell | 9 | 5 |
| Retina - Fetal Eye | 9 | 3 |
| Retina - 3D Organoid Stem Cell | 10 | 47 |
| Retina - RGC Stem Cell | 10 | 3 |
| RPE - Adult Tissue | 11 | 48 |
| Retina - Fetal Tissue | 12 | 35 |
| Brain - Amygdala | 13 | 30 |
| Brain - Anterior cingulate cortex (BA24) | 13 | 27 |
| Brain - Caudate (basal ganglia) | 13 | 28 |
| Brain - Cerebellar Hemisphere | 13 | 1 |
| Brain - Cortex | 13 | 28 |
| Brain - Frontal Cortex (BA9) | 13 | 29 |
| Brain - Hippocampus | 13 | 29 |
| Brain - Hypothalamus | 13 | 28 |
| Brain - Nucleus accumbens (basal ganglia) | 13 | 30 |
| Brain - Putamen (basal ganglia) | 13 | 28 |
| Brain - Spinal cord (cervical c-1) | 13 | 28 |
| Brain - Substantia nigra | 13 | 28 |
| Pituitary | 13 | 1 |
| Lung | 14 | 29 |
| Whole Blood | 15 | 19 |
| Brain - Cerebellar Hemisphere | 16 | 28 |
| Brain - Cerebellum | 16 | 29 |
| Brain - Cortex | 16 | 1 |
| Thyroid | 17 | 27 |
| Heart - Atrial Appendage | 18 | 30 |
| Heart - Left Ventricle | 18 | 28 |
| Skin - Not Sun Exposed (Suprapubic) | 19 | 29 |
| Skin - Sun Exposed (Lower leg) | 19 | 24 |
| Muscle - Skeletal | 20 | 29 |
| Prostate | 20 | 1 |
| Artery - Aorta | 21 | 30 |
| Artery - Coronary | 21 | 27 |
| Artery - Tibial | 21 | 28 |
| Adipose - Subcutaneous | 22 | 27 |
| Adipose - Visceral (Omentum) | 22 | 27 |
| Artery - Coronary | 22 | 3 |
| Breast - Mammary Tissue | 22 | 8 |
| Esophagus - Gastroesophageal Junction | 22 | 1 |
| Small Intestine - Terminal Ileum | 22 | 1 |
| Nerve - Tibial | 23 | 30 |
| Bladder | 24 | 4 |
| Breast - Mammary Tissue | 24 | 22 |
| Cervix - Endocervix | 24 | 2 |
| Fallopian Tube | 24 | 2 |
| Minor Salivary Gland | 24 | 23 |
| Prostate | 24 | 4 |
| Vagina | 24 | 1 |
| Pituitary | 25 | 28 |
| Kidney - Cortex | 26 | 27 |
| Stomach | 27 | 21 |
| Bladder | 28 | 5 |
| Colon - Sigmoid | 28 | 29 |
| Colon - Transverse | 28 | 29 |
| Esophagus - Gastroesophageal Junction | 28 | 28 |
| Esophagus - Muscularis | 28 | 28 |
| Small Intestine - Terminal Ileum | 28 | 27 |
| Stomach | 28 | 9 |
| Adrenal Gland | 29 | 30 |
| Small Intestine - Terminal Ileum | 30 | 1 |
| Spleen | 30 | 30 |
| Cells - Transformed fibroblasts | 31 | 30 |
| Liver | 32 | 30 |
| Cervix - Ectocervix | 33 | 2 |
| Esophagus - Mucosa | 33 | 30 |
| Minor Salivary Gland | 33 | 7 |
| Vagina | 33 | 3 |
| Cornea - Cell Line Endothelium | 34 | 9 |
| Cells - Leukemia cell line (CML) | 35 | 5 |
| Cervix - Ectocervix | 36 | 4 |
| Cervix - Endocervix | 36 | 3 |
| Fallopian Tube | 36 | 5 |
| Ovary | 36 | 4 |
| Testis | 36 | 1 |
| Uterus | 36 | 4 |
| Vagina | 36 | 1 |
| Testis | 37 | 4 |

**Supplemental Table 5:**
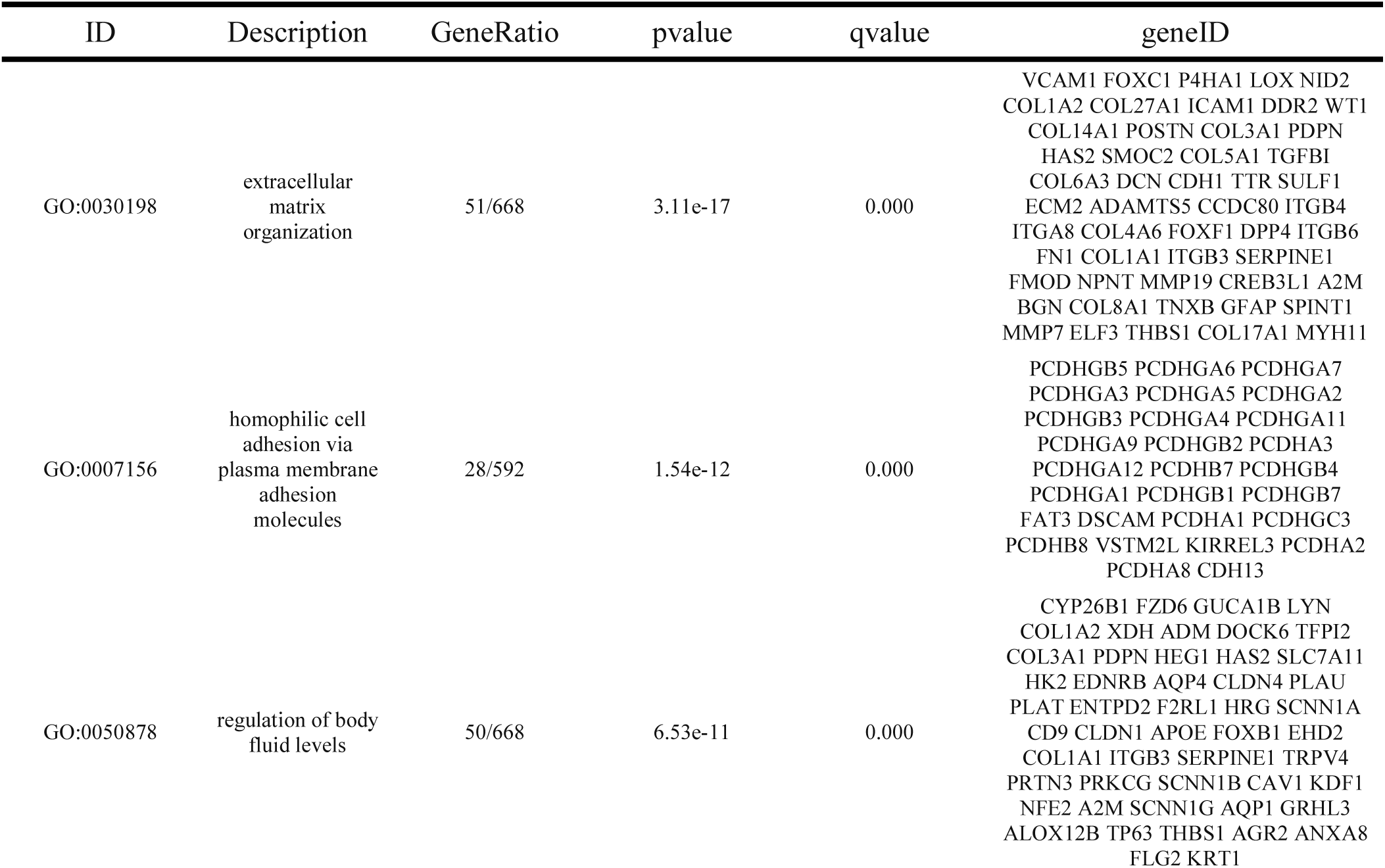
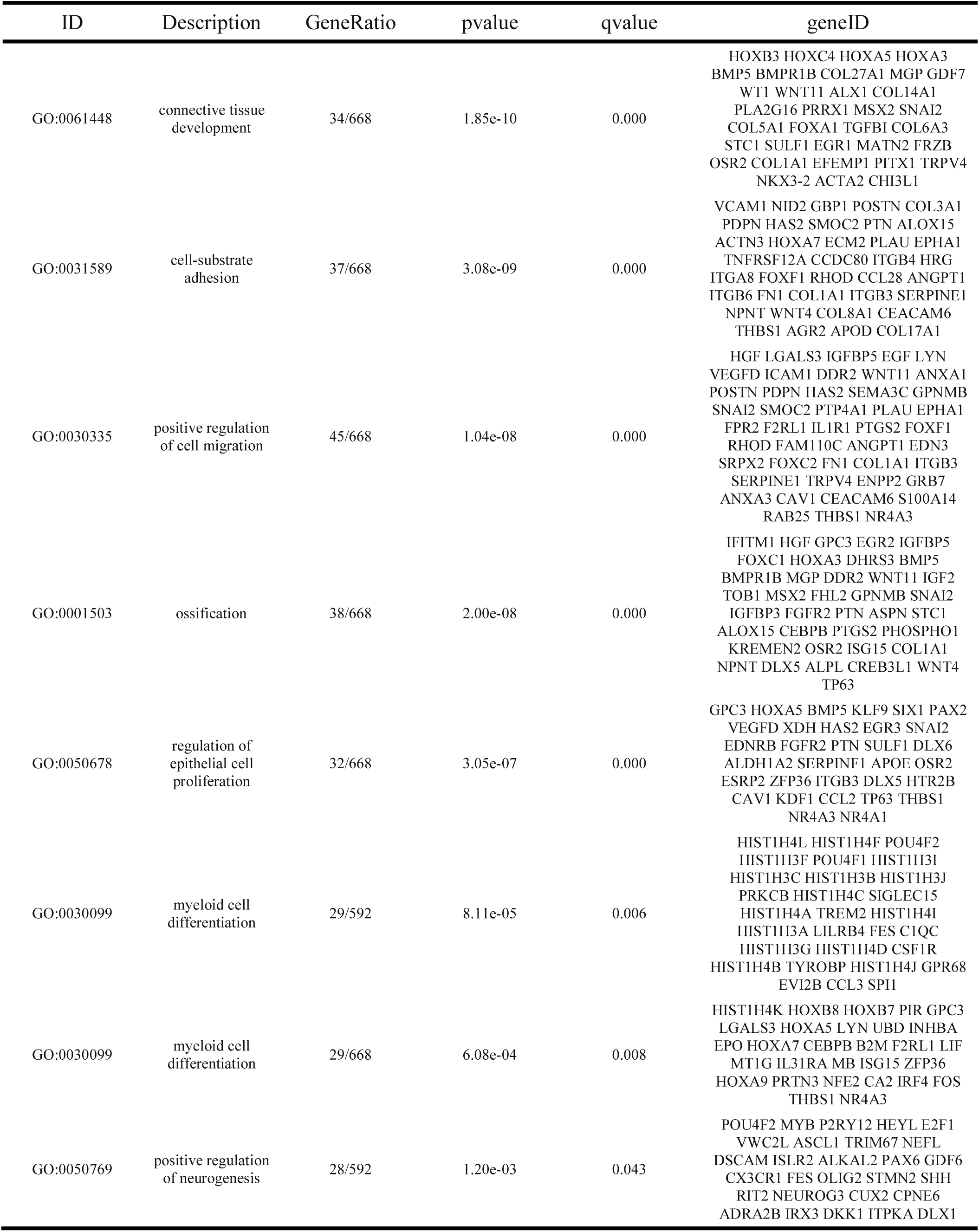
Top GO terms enriched between fetal retina and organoid retina, after paring of redundant terms with REVIGO^70^

## Acknowledgements

We would like the thank the dozens of groups who provided the raw data required to create this project. We keep a running list of the projects and associated citations at https://github.com/davemcg/eyeIntegration_app/blob/master/inst/citations.md and strongly encourage anyone who uses EiaD and eyeIntegration to cite relevant projects. We would also like to thank Kapil Bharti, Robert Hufnagel, and Brian Brooks for their continuous set of critiques and suggestions in the development of eyeIntegration app over the past two years. Tiziana Cogliati was especially helpful in the editing of this manuscript. We also would like to thank the two anonymous reviewers for their careful reading and constructive criticisms. Finally, this work utilized the computational resources of the NIH HPC Biowulf cluster (http://hpc.nih.gov).

