## Supplementary material for "Eye in a Disk: eyeIntegration human pan-eye and body transcriptome database version 1.0": power_calc.R: power_calc.R

```
# power calculation

library(ssizeRNA)
library(edgeR)
library(tidyverse)
tpm <- read_tsv('https://hpc.nih.gov/~mcgaugheyd/eyeIntegration/2019_gene_TPM_03.tsv.gz')
metadata <- read_tsv('https://hpc.nih.gov/~mcgaugheyd/eyeIntegration/2019_metadata_03.tsv.gz')

group <- metadata %>% 
  filter(sample_accession %in% names(tpm)) %>% 
  select(sample_accession, Tissue) %>% 
  unique() %>% 
  pull(Tissue)
d <- DGEList(tpm[,2:ncol(tpm)], group = group)
d <- calcNormFactors(d)
d <- estimateCommonDisp(d)
d <- estimateTagwiseDisp(d)

power <- c()
for (i in seq(5,100,5)){
  power <- c(power, check.power(nGenes = 37000, pi0 = 0.6, i, d$counts[,1], d$tagwise.dispersion, 2, up = 0.5,  replace = TRUE, fdr = 0.05, sims = 5)$pow_qvalue_ave)}

power_data <- as_tibble(cbind(seq(5,100,5), power))
save(power_data, file = 'data/power_data.Rdata')

# power_data %>% ggplot(aes(x=V1, y=power)) + geom_line() + theme_minimal() + labs(x='Number of Samples in Each Group', y = 'Power to Detect 1 log2(FC)') + scale_x_continuous(breaks = seq(10,100,10), labels = seq(10,100,10))
```
